# Genome sequencing of the nine-spined stickleback (*Pungitius pungitius*) provides insights into chromosome evolution

**DOI:** 10.1101/741751

**Authors:** Srinidhi Varadharajan, Pasi Rastas, Ari Löytynoja, Michael Matschiner, Federico C. F. Calboli, Baocheng Guo, Alexander J. Nederbragt, Kjetill S. Jakobsen, Juha Merilä

## Abstract

The Gasterostidae fish family hosts several species that are important models for eco-evolutionary, genetic and genomic research. In particular, a wealth of genetic and genomic data have been generated for the three-spined stickleback (*Gasterosteus aculeatus*), the ‘ecology’s supermodel’, while the genomic resources for the nine-spined stickleback *(Pungitius pungitius)* have remained relatively scarce. Here, we report a high-quality chromosome-level genome assembly of *P. pungitius* consisting of 5,303 contigs (N50 = 1.2 Mbp) with a total size of 521 Mbp. These contigs were mapped to 21 linkage groups using a high-density linkage map, yielding a final assembly with 98.5% BUSCO completeness. A total of 25,062 protein-coding genes were annotated, and ca. 23% of the assembly was found to consist of repetitive elements. A comprehensive analysis of repetitive elements uncovered centromeric-specific tandem repeats and provided insights into the evolution of retrotransposons. A multigene phylogenetic analysis inferred a divergence time of about 26 million years (MYA) between nine- and three-spined sticklebacks, which is far older than the commonly assumed estimate of 13 MYA. Compared to the three-spined stickleback, we identified an additional duplication of several genes in the hemoglobin cluster. Sequencing data from populations adapted to different environments indicated potential copy number variations in hemoglobin genes. Furthermore, genome-wide synteny comparisons between three- and nine-spined sticklebacks identified chromosomal rearrangements underlying the karyotypic differences between the two species. The high-quality chromosome-scale assembly of the nine-spined stickleback genome obtained with long-read sequencing technology provides a crucial resource for comparative and population genomic investigations of stickleback fishes and teleosts.

## Introduction

The small teleost fishes of the Gasterostidae family have served as important model systems in ecology and evolutionary biology, and in the study of adaptation in particular (Wootton 1976; Wootton 1984; Bell and Foster 1994; Östlund-Nilsson et al. 2006; Von Hippel 2010). The most well-known member of this family is the three-spined stickleback (*Gasterosteus aculeatus* Linnaeus, 1758), also dubbed as ecology’s and evolutionary biology’s ‘supermodel’ (Gibson 2005). The genome of the three-spined stickleback was sequenced in 2006 (Jones et al. 2012), making the species an attractive model system for studying genomic architecture of ecologically important traits, as well as for population genomic studies in general. The nine-spined stickleback (*Pungitius pungitius* Linnaeus, 1758), suggested to have diverged from the three-spined stickleback at least 13 million years ago (Bell et al. 2009), is the next most frequently utilized model species from the Gasterostidae family. It has recently gained recognition as an especially interesting model system for comparative investigations of adaptive evolution (e.g (Shapiro et al. 2006; Shikano et al. 2013; Raeymaekers et al. 2017)), sex chromosome evolution (e.g (Natri et al. 2019)) and study of adaptive divergence in the face of strong genetic drift (Merilä 2013; Karhunen et al. 2014). Although the nine- and three-spined sticklebacks share nearly identical circumpolar distribution ranges (Wootton 1984), the former shows a far greater degree of genetic population structuring than the latter (Shikano et al. 2010; DeFaveri et al. 2011). This, together with the fact that the genus *Pungitius* appears to be more specious than the genus *Gasterosteus* (8-10 *vs.* 3 species, respectively; Eschmeyer 2015), suggests that there are likely important differences in processes and forces governing differentiation between each of the two species. Hence, this species pair provides an interesting model system for comparative and population genomic investigations aiming to disentangle the relative importance of factors influencing processes of population differentiation and speciation.

While genomic resources for the three-spined stickleback, including an annotated reference genome (Jones et al. 2012), are well-developed, those for the nine-spined stickleback are rather less developed, typically relying on the three-spined stickleback reference genome. The recently published ultra-high density linkage map (Rastas et al. 2015; Li et al. 2018) and a draft version of the nine-spined stickleback genome based on short-read sequencing technology (Nelson and Cresko 2018) are important developments in this regard. However, short-read technology based draft assemblies are often of limited utility when dealing with fairly large and complex vertebrate genomes. High-contiguity chromosome-level assemblies obtained from long-range information are vital for resolving large repetitive regions and providing robust insights into genome and chromosome evolution. Therefore, a high-quality genome assembly based on long-read technologies for the nine-spined stickleback would provide a valuable resource for comparative and population genomic studies of stickleback fishes.

Furthermore, a particularly high degree of divergence in karyotype characterized by varied chromosomal morphology and diploid numbers in sticklebacks has been previously noted (R. Chen and Reisman 1970; Ocalewicz et al. 2011; Urton et al. 2011). The three- and nine-spined sticklebacks have a diploid chromosome number (2n) of 42, and the number of arms (NF) of 58 and 70 respectively, while the four-spined (*Apeltes quadracus*) and brook stickleback (*Culaea inconstans*) karyotypes have 23 pairs of chromosomes. Hence, the 2n of nine-spined stickleback is more similar to that of three-spined stickleback than to those of the more closely related four-spined and brook sticklebacks. Thus, the exact ancestral karyotype of sticklebacks is not well understood in relation to the phylogeny (Kawahara et al. 2009) of the family.

Here, we present the first chromosome-level genome assembly of the nine-spined stickleback. The high-coverage long read PacBio sequencing integrated with an ultra-dense linkage map yielded a high-quality contiguous assembly ordered into 21 pseudo-chromosomes. Using this new resource, we provide a comprehensive analysis of repetitive elements including centromeric repeats in the nine-spined stickleback genome. We also describe a recent duplication in the MN cluster of hemoglobin and show that this region could potentially involve frequent CNVs in the species. Utilizing our chromosome-scale assembly, we now identify and pin-point structural variations potentially explaining the divergent karyotypes of the three- and nine-spined sticklebacks.

## Results

### Genome assembly and validation

We sequenced a male *P. pungitius* individual using the PacBio RSII platform yielding ~110x of genome coverage. The error-corrected reads were assembled using Celera assembler (Miller et al. 2008) followed by polishing with Quiver (Chin et al. 2013). The assembly improvement was done by mapping Illumina HiSeq 2500 reads to the polished assembly using Pilon (Miller et al. 2008; Walker et al. 2014). This resulted in an initial assembly consisting of 5,305 contigs with a total size of 521 Mb and an N50 of 1.2 Mb (Table 1). The total assembly size is close to the genome size estimates of about 550-650 Mbp in other stickleback species (Hinegardner and Rosen 1972; Vinogradov 1998).

**Table 1.** Assembly statistics.

|  | **Raw assembly** | **Anchored assembly** |
| --- | --- | --- |
| Number of contigs | 5303 | 4939* |
| Total size of contigs | 521,182,237 | 521,203,469 |
| Longest contigs | 9,719,887 | 32,096,348 |
| N50 scaffold | 1,233,545 | 17,578,551 |
| **Assembly Validation** |  |  |
| Complete BUSCOs | 97.1 % |  |
| Complete Single-copy BUSCOs | 4269 (93%) |  |
| Complete Duplicated BUSCOs | 183 (4%) |  |
| Fragmented BUSCOs | 62 (1.4%) |  |
| Missing BUSCOs | 68 (1.5%) |  |
| Total BUSCO groups searched | 4584 |  |

\* 21 LGs and unplaced contigs

A high-density linkage map of almost 90,000 markers and 1,000 individuals was constructed and used to order and orient the majority of the assembled contigs. Utilizing this map, we placed 686 contigs, comprising ~444 Mb (~85%) of the assembly, into 21 pseudo-chromosomes. The largest assembled pseudo-chromosome LG12, which is also the sex chromosome of Eastern European lineage of *P. pungitius* (Shapiro et al. 2009; Rastas et al. 2015; Natri et al. 2019), is of size ~40.9 Mbp. To validate the placement of the contigs, we inspected the collinearity between the physical and linkage maps. A high degree of correspondence was observed between the marker order in the linkage map and the assembled pseudo-chromosomes (*r* = 0.95; Figure 1). Consistent with expectations, there was a general monotonic increase between the physical and genetic maps along most of the pseudo-chromosomes, except for some regions corresponding to lower recombination rates (Figure 1). Further validation of the assembly was performed by assessing the genome completeness using BUSCO (Simão et al. 2015). The results indicated high contiguity with 98.5 % of BUSCO genes found complete or fragmented in the assembly (Table 1 and Figure S1).

**Fig. 1.**
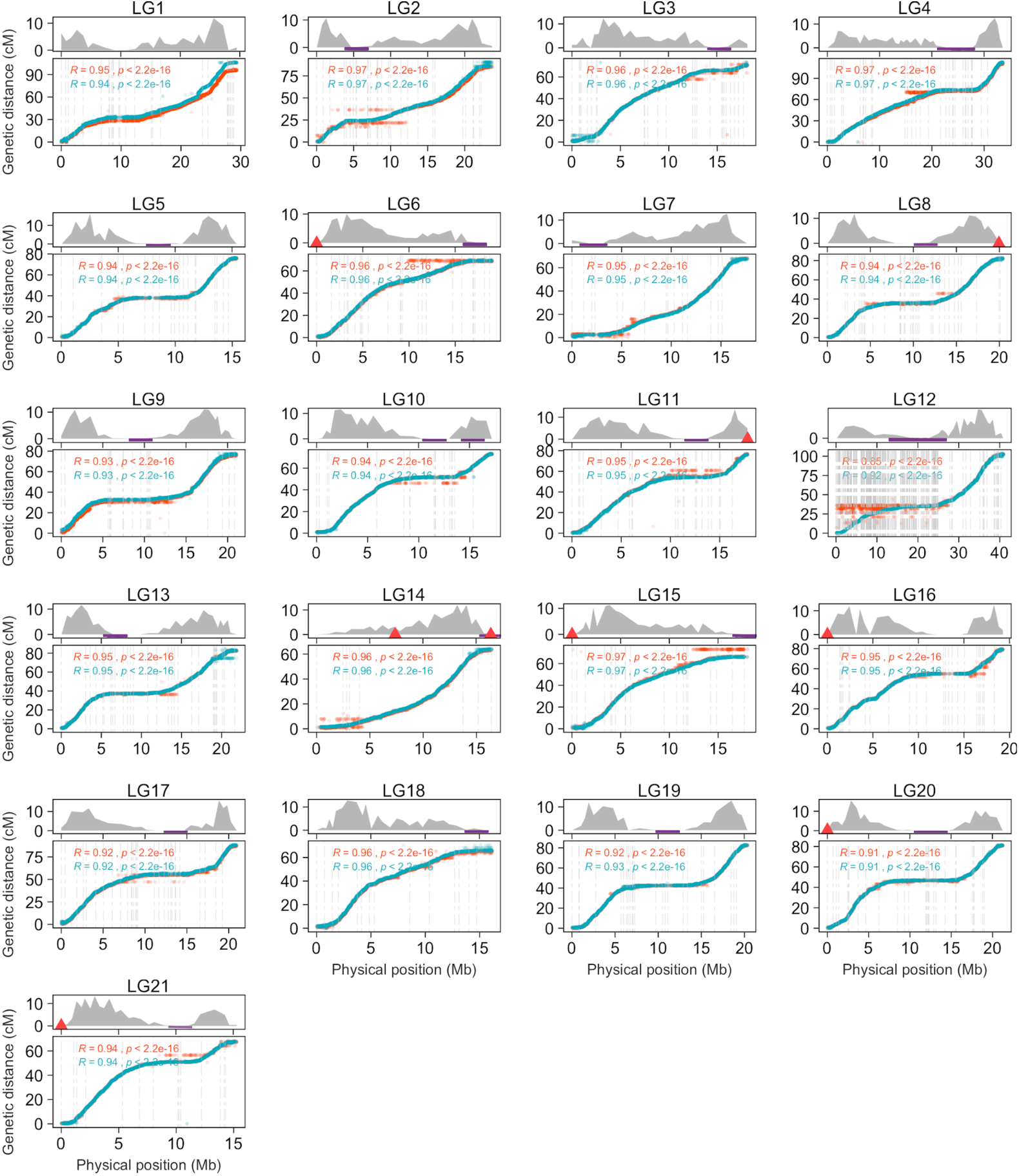
Concordance of the assembly with linkage map. The plots represent the correspondence between genetic (cM; Y-axis) and physical (Mb; X-axis) positions of the markers on each of the pseudo-chromosomes (bottom panels). The turquoise points correspond to the sex averaged map. The orange points are map positions from a technical replicate using a different subset of markers (see Methods). The dashed lines represent the contig borders and the maximum value on x-axis corresponds to the size of the pseudo-chromosomes in the assembly. The corresponding recombination rates (cM/Mb) are plotted along the length of each pseudo-chromosome (top panel). The potential telomeric regions in the assembly are marked with red triangles and the purple rectangles represent locations of the identified centromere-associated tandem repeat in the nine-spined stickleback genome.

### Genome annotation

We constructed a nine-spined stickleback-specific repeat library using *de novo* and homology-based approaches (see Methods). Redundant sequences were removed, and the remaining sequences were classified (see Methods). Sequences with hits to Uniprot/SwissProt database (UniProt Consortium 2015) were removed and the remaining 1,450 sequences were then combined with teleost repeat sequences from Repbase (Bao et al. 2015). Repeat masking using the custom-made repeat library identified 23.16% of the genome assembly as repetitive. A combination of evidence-based and *ab initio* gene predictions followed by filtering based on ‘annotation edit distance’ (AED) score and presence of PFAM domains resulted in 25,062 high confidence gene models. Of these, 22,925 reside on the pseudo-chromosomes and the remaining 2,137 on unplaced contigs.

### Genome-wide characterization of transposable elements

Analysis of transposable elements (TEs) in the nine-spined stickleback assembly revealed that the genome comprises 6.91% of DNA transposons, ~4.60% of LTR retrotransposons and 2.28% of LINE and 0.5% of SINE elements (Figure 2a). To facilitate comparison, we created a three-spined stickleback-specific repeat library using the same approach, utilizing the genome assembly generated in Glazer et al (2015). Using this set of repeats, we classified 16.22% of the three-spined stickleback genome as repetitive, with DNA transposons accounting for 3.73%, LTRs for 3.13% and LINE and SINE elements comprising of 2.76% and 0.32% respectively. This estimate for three-spined stickleback is close to that obtained in other studies (e.g. (Gao et al. 2016), (Chalopin et al. 2015)). Similar to earlier observations, the three-spined stickleback genome was found to have no particular predominance of transposable element families and a similar lack of dominance was observed for the nine-spined stickleback, albeit DNA transposons were slightly more abundant than LTR elements. Further, the diversity of the repeat families is fairly similar in the two species, while proportions vary. The abundance of different categories of repetitive elements in the two species are shown in Figure 2a. The assembly quality and possible collapse of repetitive sequences naturally affect repeat annotation and thus the overall lower proportion of repeats in the three-spined stickleback genome is likely an underestimation. We also note the general lack of accumulation of repeat families in the nine-spined stickleback genome, supported by the observation that most repeat families seem to have recent activity (Figure 2c). This is consistent with a previous study of non-LTR retrotransposons in the three-spined stickleback and suggest that active DNA loss leads to lower accumulation (Blass et al. 2012).

**Fig. 2:**
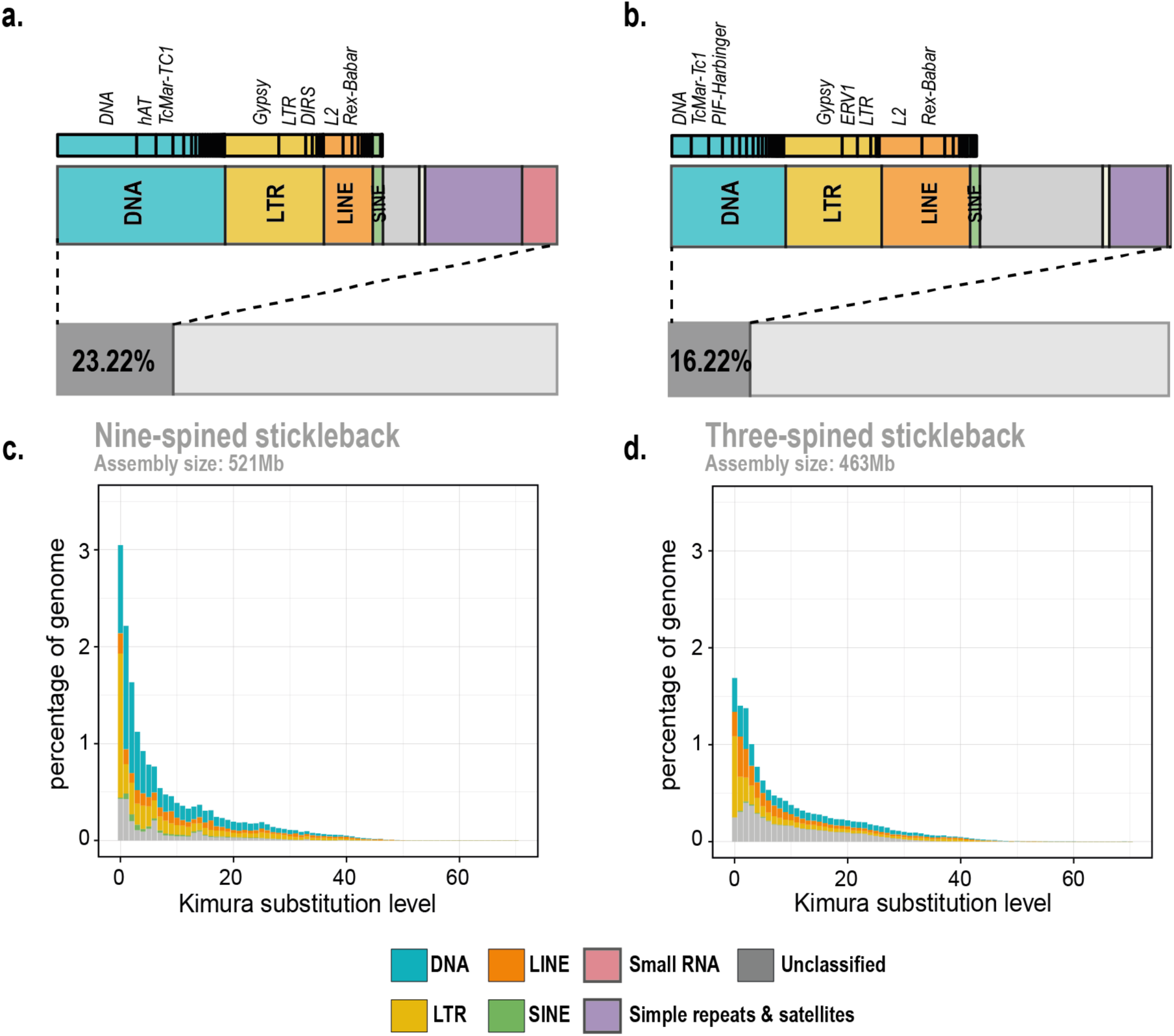
Transposable elements in stickleback genomes. **a** and **b** represent the fraction of the assembly comprising of repetitive elements in nine- and three-spined stickleback (Glazer et al. 2015) assemblies, respectively. Repeat landscapes representing the divergence of the repeat sequences to the consensus are represented for nine- and three-spined stickleback genomes in **c** and **d** respectively.

To study the activity of transposable elements, we estimated the sequence divergence between the repeat copies as a proxy for their age. We found that LTR are enriched in the youngest category and thus seem to have been recently active. In small populations, selection against repeats and elimination of accumulated repeat copies loses power and active repeat families may rapidly expand in size (Lynch and Conery 2003). To look for such patterns in the reference individual and the respective population, we sequenced (to 10X coverage) five additional individuals from the same small pond population, Pyöreälampi, Finland (FIN-PYO), and five individuals from a large marine population, Levin Navolok Bay, Russia (RUS-LEV), from which the pond population has, most probably, been derived (Supplementary figure S2). We found that, with the exception of LINEs and Tc1 DNA elements, all repeat families have significantly higher mean normalized read depths in the pond individuals than in the marine individuals. The most notable differences, 24.0 and 31.1% increases, were seen in LTRs and its Gypsy subfamily (one-sided Welch t-test: p=6.0e-06 and p=3.0e-06). The mean normalized coverage for other repeat families was 0.903–1.087, close to the expected coverage of one and demonstrating that the assembly represents the true number of repeats well, whereas the coverage for LTRs, and specifically Gypsy elements, were 1.804 and 2.219, respectively. While these numbers indicate recent activity of the particular repeat families in the genome, they might also thus be underrepresented in the assembly.

### Characterization of short tandem repeats

Short tandem repeats (STRs) are ubiquitous elements comprising of tandemly repeated units of length 1-6 bp in length (Tóth et al. 2000). STRs are implicated in various facets of genome evolution like gene regulation, chromatin organization and so on (Kashi et al. 1997; Li et al. 2002). Although a major portion of these repeats are known to reside in the non-coding intronic and intergenic regions of eukaryotic genomes, certain types of STRs have been known to occur in coding regions. The STRs in coding regions are predominant tri-nucleotide (multiples of three) repeats owing to the strong selection to maintain the reading frame (Metzgar et al. 2000). We surveyed the STR abundances in the nine-spined stickleback genome using Phobos. STRs of lower unit size appeared to be more prevalent than those with larger unit sizes, with the dinucleotide repeats being overall the most abundant class of STRs in the assembly (Supplementary Figure S3a). The proportions of STRs of different motif sizes were fairly identical with slight variation among the pseudo-chromosomes (Supplementary Figure S3b). However, the non-random distribution in different genomic regions is apparent with the overall relative abundance of STRs in intronic and intergenic regions being many folds higher than in genes (Supplementary Figure S3c). As expected, the repeats of unit size three, six and nine were relatively more abundant in the genic regions while the dinucleotide repeats were far more abundant in the non-coding regions (Supplementary Figure S3c). Among the dinucleotide repeats, ‘AC’ was the most abundant motif, followed by ‘AG’. ‘AGG’ was the most abundant triplet repeat motif across all regions (Supplementary Figure 4). In exonic regions, this was followed by the ‘AGC’ and ‘CCG’ motifs, both of which were relatively underrepresented in the intronic/intergenic regions. In addition, the ‘AAT’ motif was fairly abundant in intergenic and intronic regions but rare in the coding regions (Supplementary Figure 4). These findings are in agreement with observations from other eukaryotic genomes (Stallings 1994; Tóth et al. 2000; Li et al. 2004). Additionally, we also looked for telomeric tandem repeats characterized by a typical conserved G-rich hexamer motif ‘TTAGGG’. Large telomeric arrays were found in LG8, LG11, LG14 and LG21 comprising 975.33, 272.17, 1,663 and 1,033.17 copies of the telomeric repeats respectively (marked in Figure 1).

### Characterizing centromeric repeats in the nine-spined stickleback genome

A substantial portion of eukaryotic genomes is comprised of satellite repeats. Large satellite tandem repeat arrays are often observed in the heterochromatin, including centromeric and pericentromeric regions, and frequently constitute the most abundant tandem repeat category in genome assemblies (Melters et al. 2013). Although the exact role of these repeat elements in structure and function of centromeres is not well understood, they are thought to be vital for various processes such as chromosome segregation, proper pairing of homologous chromosome and packaging of centromeric DNA (Plohl et al. 2008). These highly repetitive structures of heterochromatin are still a major impediment to proper genome assembly and mapping of such regions. Our long-read-based assembly allows insights into the sequence composition of these regions. Using the nine-spined stickleback-specific repeat library, ~18.4% of the assembly was annotated as known repetitive elements and the rest remained unclassified. The predominant repeat among the unclassified sequences occupied up to ~2.2% of the assembly, and accounted for about 45% of the bases masked by the unclassified repeats in the library. This repeat consisted of tandem repeat units with sizes of 176-180 and ~360 bp. The low GC content of the sequence along with the distinct monomer size, often associated with centromeric satellites, warranted further analysis. To characterize the centromeric repeat monomer size and abundance, we performed a search for tandem repeats on a randomly selected subset of ~500,000 PacBio subreads using Tandem Repeat Finder (Benson 1999). The results were parsed using a method similar to that used by Melters et al. (Melters et al. 2013). Specifically, we retained only the shortest monomer representing the repeats longer than 50 bp and covering a minimum of 80% of the read length. The resulting repeats show a clear peak around a monomer length of ~178 bp (Figure 3a). Further, the AT-rich 178 bp monomer is organized into dimer (~350 bp) and trimer (~530 bp), as apparent from the distinct peaks (Figure 3a).

**Fig. 3:**
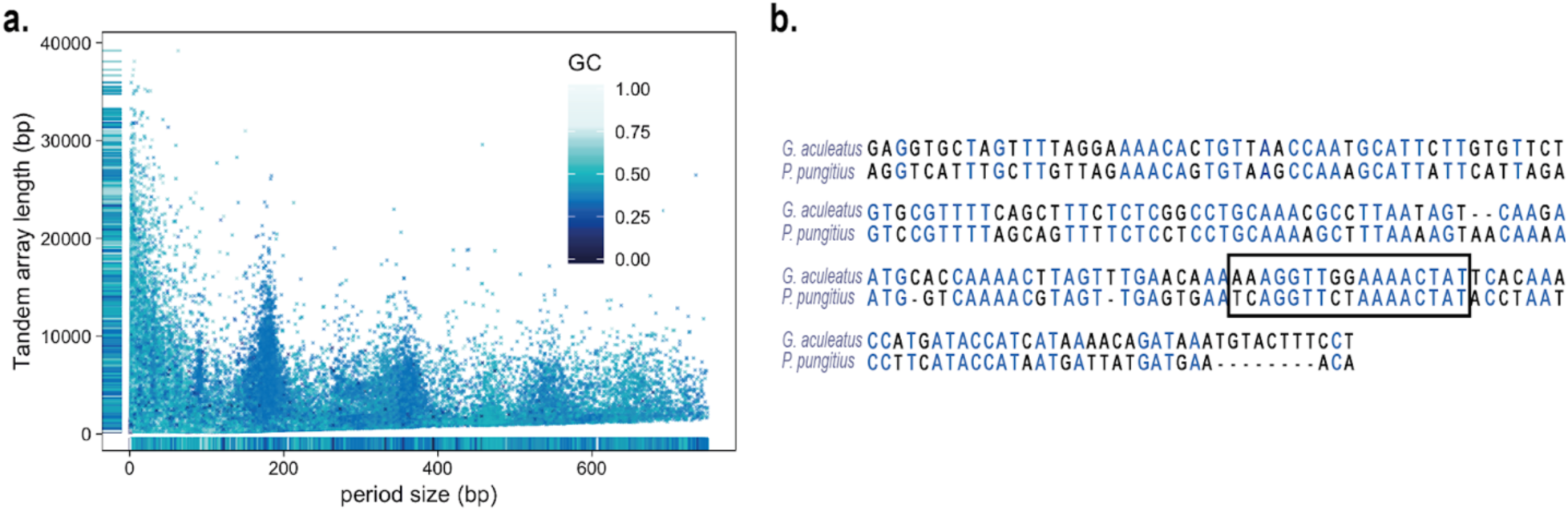
Distribution of tandem repeats in the nine-spined stickleback assembly. **a.** The x-axis represents period size of the tandem repeat units and the y-axis represents array size (period size x number of repeats). The points are colored based on GC content **b.** Alignment of the nine-spined stickleback putative centromeric repeat to the centromeric repeat sequence in the three-spined stickleback (GacCEN, accession KT321856 (Cech and Peichel 2015); identity 61.2%). Identical bases are represented in blue. The CENP-B box in three-spined stickleback is marked with a dashed rectangle.

Although functionally conserved, owing to the rapid evolution of the satellite DNA, there is often a significant sequence divergence in centromeric repeats among related species, with sequence similarity detectable only in species that have diverged within ~50 MYA (Melters et al. 2013). In three-spined stickleback, an AT-rich 186 bp repeat was identified and confirmed to occur in centromeric constrictions (Cech and Peichel 2015). The alignment of the centromere-associated repeat sequence in nine-spined stickleback to that the three-spined stickleback centromeric repeat (GacCEN), showed a considerable sequence similarity (~61%) and in the CENP-B box region in particular (Figure 3b).

Further, one of the distinguishing features of centromeric and pericentromeric regions is the apparent suppression of recombination. Thus, we investigated the positions of the representative centromeric repeat relative to the recombination rates along the pseudo-chromosomes. The identified repeat indeed consistently corresponded to regions of low recombination, marking pericentromeric regions (Figure 1, top panels).

To understand how various genomic features vary relative to each other, we analysed the distribution of GC content, gene density, transposable elements and recombination rate along the assembled pseudo-chromosomes. In line with expectations, GC content and gene density were generally reduced in areas of low recombination, while TEs were enriched. To access the global trend of this variation, we computed correlations of recombination rates with GC, gene density and repeat density (Supplementary Figure S5 and S6). Indeed, GC content shows a significant positive correlation with recombination rate (p-value<2.2e-16) and transposable element density shows a strong negative correlation (p-value<2.2e-16), whereas the gene density is only weakly positively correlated with the recombination rate (p-value=4.1e-05). Interestingly, TE density also shows a negative correlation with the density of STR density (p-value<2.2e-16). Further, the approximate pericentromeric region for each of the pseudo-chromosomes was defined based on the location of the inferred centromeric tandem repeat and reduction of recombination rates (for LG1 and LG16, we used only the region of low recombination). Using these compartments for comparison, we found a significant increase in GC content and gene density outside pericentromeric regions and a significant enrichment of TEs in the pericentromeric regions (Supplementary Figure S6). Apart from the general enrichment of TEs in the pericentromeric regions, we compared the relative proportions of LTR and DNA elements to the total TE content for each bin across each of the pseudo-chromosomes to look for enrichment of specific classes of TEs. Overall, LTRs, specifically gypsy elements, are consistently more enriched in pericentromeric regions than outside of them (Wilcoxon p-value<2.2e-16) and thus are likely associated with centromeric regions in the nine-spined stickleback chromosomes (Figure 4). Next, we defined pericentromeric region in the three-spined stickleback chromosomes using locations of the GacCEN repeat (Cech and Peichel 2015) and including a 2 Mb flanking region on either side of the repeat. A similar increase in relative proportions of LTR elements was observed in the pericentromeric regions of the three-spined stickleback chromosomes (Supplementary Figure S7).

**Fig. 4:**
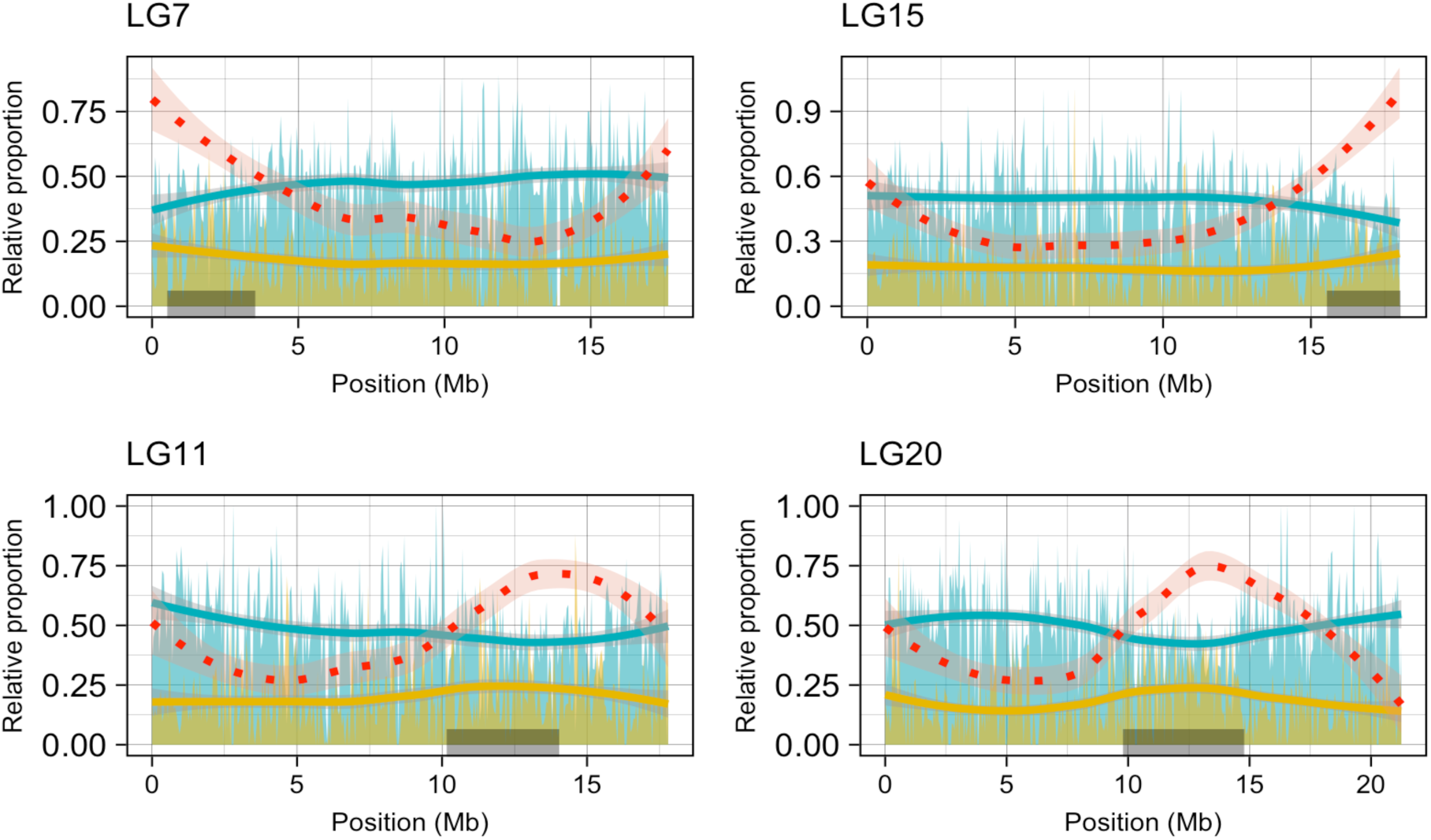
Examples of relative proportions of LTR and DNA transposons along nine-spined stickleback pseudochromosomes. Distribution of relative proportions of LTR (yellow) and DNA TEs (blue) transposable elements to total repeat content per 50Kb bin along selected pseudo-chromosomes (LG7, 11, 15 and 20). The red dotted line represents log10 of absolute abundance of LTR-gypsy elements across the pseudo-chromosomes. The gray rectangles (at the bottom) depict the pericentromeric region.

### Gene-based phylogeny and gene family evolution

To assess global gene content evolution, we compared the inferred protein-coding gene set with proteins from eight other teleost species including zebrafish, Atlantic cod, platyfish, medaka, fugu and three-spined stickleback. A total of 17,976 orthogroups were inferred using Orthofinder (Emms and Kelly 2015), of which 9,811 are present in all nine species and 5,098 were single-copy orthologs in the nine species. The two stickleback species share a set of 71 core orthogroups.

To infer the divergence times among the nine species, we retained 2,691 single-copy orthologs that were also a part of BUSCO Actinopterygii (odb9) genes set. The resulting sequences were concatenated into a supermatrix of 2,085,162 bp that was used for phylogenetic reconstruction and divergence time estimation with BEAST2 (Bouckaert et al. 2014). The phylogeny of the nine species was time calibrated by placing age constraints according to the timeline of teleost evolution inferred by Betancur-R. et al. (2013). These age constraints were placed on all internal nodes except the divergence of the two stickleback species. Using this approach, the divergence time between nine- and three-spined sticklebacks was estimated to be around 27.1 million years ago (MYA), with a 95% highest posterior density interval (HPD) of 25.5 - 28.8 MYA. To assess the robustness of the estimate, we performed separate BEAST analyses for each of the 2,691 individual gene alignments. This resulted in a distribution of divergence time estimates with the median and mean of 23.0 and 26.6 MYA, respectively.

Although the BUSCO genes should appear as single copy in all species, the full data set may contain non-homologous sequences. To reduce the error from these, we applied a set of rigorous filters. First, we removed 24 gene trees that did not support monophyly of the three- and nine-spined sticklebacks. These genes were excluded from further analyses because, given the long timescales separating sticklebacks from other species included in the phylogeny, their apparent non-monophyly is more likely to result from low phylogenetic signal than from processes like incomplete lineage sorting. Second, to avoid errors due to paralogy and misalignments, we performed stringent filtering of the protein alignments generated by Orthofinder and retained only the 825 high-confidence orthogroups (see methods). Finally, we discarded genes not evolving in a clock-like manner. By selecting the remaining genes conforming, at least to some extent, to the molecular-clock assumption (Zuckerkandl and Pauling 1962), greater precision of divergence time estimates can be expected. In addition, as unidentified paralog sequences would likely increase the inferred rate variation, the selection of clock-like genes also further reduces the probability of paralogs remaining in the alignments. The above filtering steps resulted in a total of 778 orthogroups and a concatenated alignment of 548,248 bp. This concatenated alignment was analyzed with BEAST under the same settings as the full supermatrix. Using the concatenated data set, the divergence time between nine- and three-spined stickleback was estimated to be around 25.8 MYA (95% HPD 18.9-35.04 MYA, Figure 5a) whereas the separate analysis of 778 gene alignments gave median and mean divergence time estimates of 23.1 and ~25.7 MYA, respectively (Supplementary Figure S8a). To assess the impact of the calibration points, we repeated the analysis of 778 gene alignments with only two calibration points (indicated by green asterisks in Figure 5a). The median estimate from this analysis was 23.9 MYA (mean of 27.1 MYA, Supplementary Figure S8b).

**Fig. 5:**
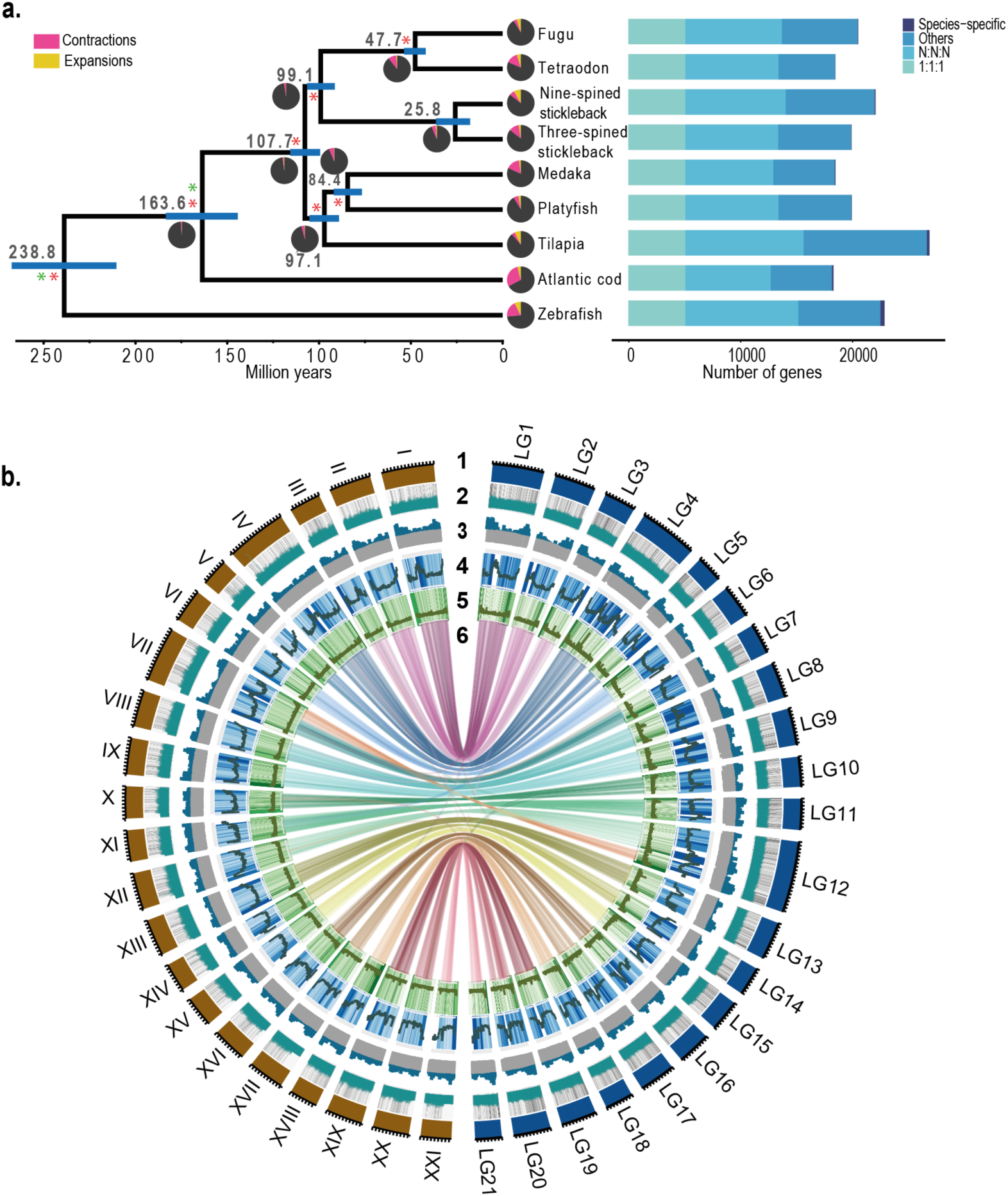
Evolutionary and comparative genomic analysis. **a.** Phylogenetic tree using orthogroups inferred from nine teleost species. The number of gene families expanded and contracted have been indicated in the pie diagrams in red and yellow, respectively. **b.** Circos plot representing gene-level synteny between the nine-(right) and three-spined stickleback (left) genome assemblies. Tracks: 1) nine- and three-spined chromosomes, 2: gene density, 3: GC content, 4: TE density, 5: tandem repeat density and 6: links of synteny (defined by 10 collinear genes) between the two species.

### Identification of a recent tandem duplication in the hemoglobin MN cluster

Synteny analysis using MCScanX (represented as ribbons in Figure 5b) was further utilized to identify tandem duplicates across the assembly. Interestingly, one such duplication was indicated in the hemoglobin cluster in LG11. We therefore investigated the hemoglobin repertoire in nine-spined stickleback, using alpha and beta globin genes from three-spined stickleback and zebrafish as query sequences. In line with the expectation from three-spined stickleback genome, the MN and LA clusters were found in LG11 and LG5 of the nine-spined stickleback assembly. While the arrangement of alpha (Hba) and beta (Hbb) hemoglobin genes on opposite strands in MN cluster is conserved between the species, we found four more alpha and beta globin genes (leading to 15 in total) in the nine-spined than in the three-spined stickleback genome. The entire set of genes was present in a single contig of the raw nine-spined stickleback assembly. To investigate for lineage-specific duplications in the nine-spined stickleback genome, we firstly excluded the possibility of misassembly by mapping and computing the depth of Illumina reads across the genes. We then performed a self-alignment of the hemoglobin cluster in the nine-spined stickleback and found high internal similarity in the region, extending outside exons and into intergenic regions (Figure 6a). This pattern suggests recent, *en bloc* duplications likely comprising of multiple Hba and Hbb genes. To estimate the age of the gene duplications, we inferred a phylogenetic tree from the predicted protein sequences (Supplementary Figure S9). The genes comprising the putative duplication block form a distinct clade next to the syntenic genes in the three-spined stickleback. As the high sequence identity indicates a very recent duplication event, we decided to study the region in more detail using population genomic data. For this, we used the data from five sequenced individuals from both the Pyöreälampi pond (Finland) population and the ancestral marine population from Levin Navolok Bay, the Baltic Sea (Russia). The normalised mean sequencing coverages for individual hemoglobin genes show that the duplication region is fixed in the pond but missing in the ancestral marine population (Figure 6b). This suggests that the duplication has happened since the split of the two populations at most 8000 years ago, after the retreat of the ice sheet in North-eastern Fennoscandia (Shikano et al. 2010; Bruneaux et al. 2013; Wang et al. 2015). Interestingly, other genes within the same cluster show higher coverage in the marine population and thus additional duplications of individual hemoglobin genes may be relatively frequent.

**Fig. 6:**
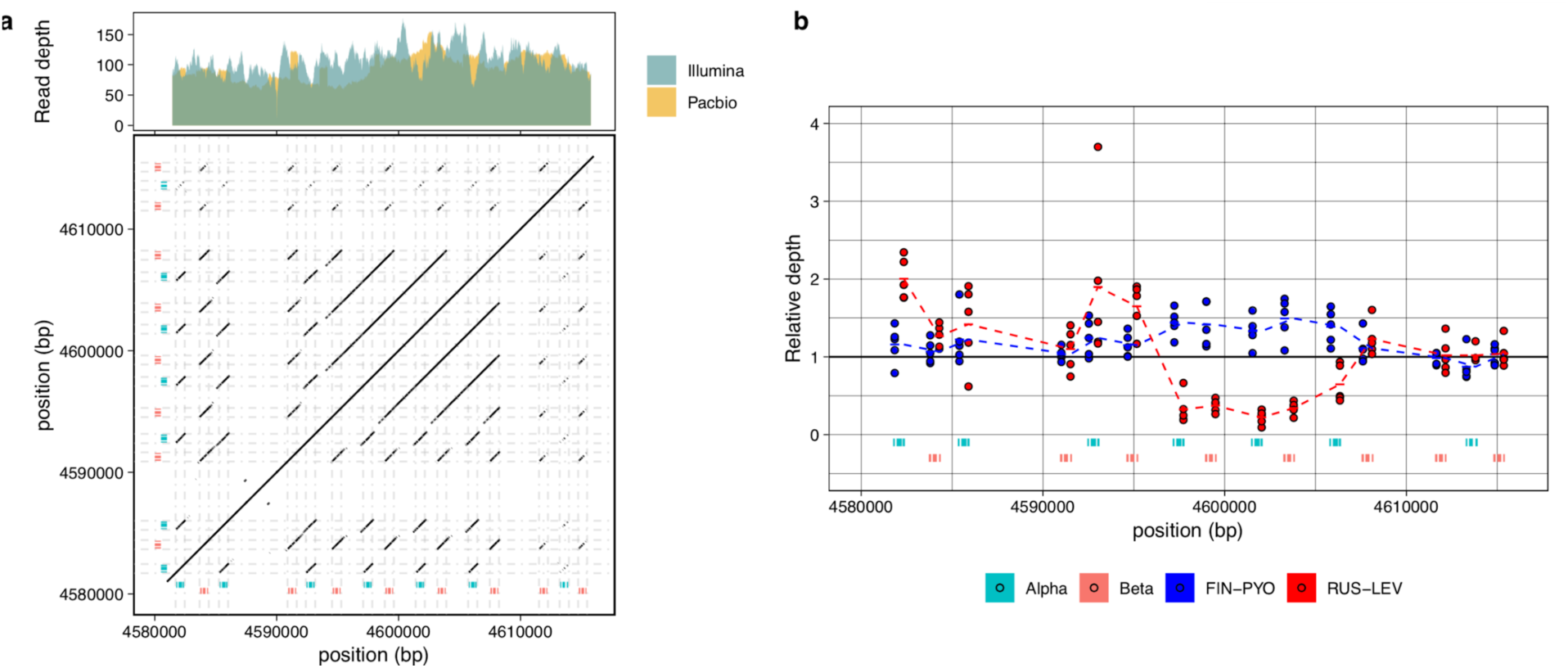
Analysis of hemoglobin MN cluster. **a.** Dotplot (bottom) representing self-alignment of the MN cluster region of the hemoglobin in nine-spined stickleback. Top: Mapped read depth across the MN cluster of hemoglobin in nine-spined stickleback. **b.** Relative read depth for five individuals each from nine-spined stickleback populations FIN-PYO (Pyöreälampi pond, Finland) and RUS-LEV (Levin Navolok Bay, Russia). The dashed lines connect the means of the read depth for each of the hemoglobin genes.

**Fig. 7:**
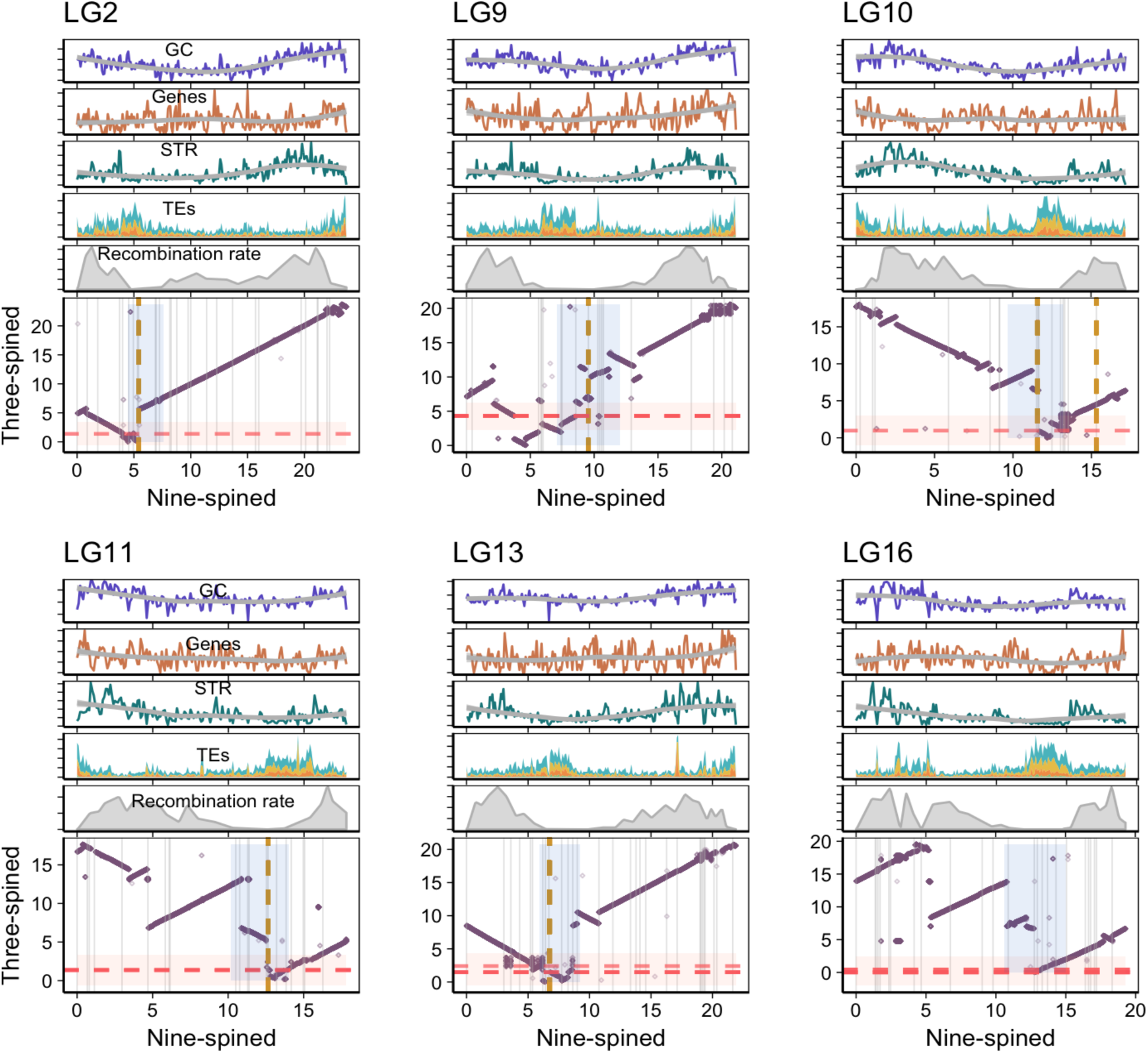
Conserved synteny between three-spined and nine-spined stickleback for chromosomes (LG2, 9, 10, 11, 13 and 16) potentially involving inversions that have led to divergent chromosomal morphology between three-and nine-spined sticklebacks. Top: The distribution (per 100 kb bins) of GC content, gene density, TE density (yellow: LTR, blue: DNA, orange: LINE, green: SINE) and recombination rate along chromosome 11. Bottom: Alignment of nine-(x-axis) and three-spined (y-axis) stickleback chromosomes. The shaded areas represent the putative pericentromeric region in the two genomes.

### Chromosome evolution in sticklebacks

While most teleosts have a karyotype comprising 24 pairs of chromosomes, three fusion events involving chromosomes 1, 4 and 7 were found to likely result in the reduced karyotype of the three-spined stickleback (Amores et al. 2014). Based on our macrosynteny analysis, the fusion of chromosomes 1 and 4 is distinctly detected in the nine-spined stickleback genome with 1:1 synteny correspondence across chromosome arms when compared with the corresponding ortholog in medaka (Supplementary Figure S10). Generally, a high degree of overall genomic collinearity is observed between the nine- and three-spined stickleback chromosomes (Figure 5b). However, a considerable number of intra-chromosomal rearrangements were observed. Such a lack of gene order conservation was earlier described by comparing linkage maps between the two species (Rastas et al. 2015). The translocation of the three-spined stickleback chromosome 7 arm to the sex chromosome (LG12) of nine-spined stickleback (Shikano et al. 2013; Rastas 2017) was the only major inter-chromosomal rearrangements observed.

Both 2n and NF (number of chromosome arms) vary considerably across sticklebacks, with nine-spined stickleback having a 2n of 42 with highest NF of 70 among stickleback species with a distinctly larger number of metacentric and submetacentric chromosomes than the three-spined stickleback (Ocalewicz et al. 2011). A comparison of cytogenetic data of the four- and three-spined sticklebacks revealed various rearrangements leading to the differences in NF between the two stickleback species (Urton et al. 2011). A common mechanism to achieve an increase in NF is by pericentric inversions, as these retain 2n while increasing NF. Between the three-spined and four-spined sticklebacks, six pericentric inversions, involving chromosomes 1, 3, 8, 17, 20 and 21, have been observed (Urton et al. 2011). Firstly, we attempted to investigate the synteny relationships within these chromosomes to infer potential rearrangements in the nine-spined stickleback chromosomes. The pericentric inversion of chromosome 1 (acrocentric in four-spined stickleback and metacentric in three-spined stickleback) seems specific to the four-spined stickleback karyotype since the chromosome 1 seems to be syntenic along the entire length (and metacentric) between three- and nine-spined sticklebacks. The pericentromeric inversions on the chromosomes orthologous to three-spined stickleback chromosomes 8 and 20 are likely shared by four-spined and nine-spined sticklebacks, both of which seem metacentric, thus reflecting likely rearrangements in the common ancestor of four- and nine-spined sticklebacks (Supplementary Figure S11). We also observe a pericentric inversion in LG21, while the rearrangements in LG17 and LG3 are rather unclear (Supplementary Figure S11). Since the nine-spined stickleback karyotype has a larger number of bi-armed chromosomes than that of the three-spined stickleback, we looked for additional pericentromeric inversions explaining this difference. First, we inferred karyotypes based on the pericentromeric regions defined in the assembly and looked for chromosomes that differ in morphology between three- and nine-spined sticklebacks. Indeed, we find that LG2, LG10, LG11, LG13 and LG16 of the nine-spined stickleback harbor potential NF-increasing inversions leading to different chromosomal morphologies in the two species (Figure 6). We further confirmed these rearrangements based on synteny with corresponding medaka chromosomes (Supplementary Figure S12).

## Discussion

We have generated and described a high-quality chromosome-level genome assembly of the nine-spined stickleback. The raw assembly using PacBio long reads yielded 5,305 contigs and 686 of these were anchored into 21 pseudochromosomes with the aid of a high-resolution linkage map, representing 85% of the assembly size. About one-fourth of the nine-spined stickleback genome was found to consist of repetitive sequences. The completeness of the assembly enabled in-depth investigation of these highly repetitive regions of the genome.

In eukaryotic chromosomes, centromeric heterochromatin is often known to be dominated by satellite DNA consisting of large homogenized tandemly arranged arrays of monomeric repeat sequences (Melters et al. 2013). Our survey of tandem repeats in the assembly identified a high-copy 178 bp long centromere-associated repeat organized in such large arrays. This length of identified repeat is consistent with the length of 150-180 bp required to wrap around a single nucleosome unit (Henikoff 2001).

Furthermore, transposable elements were found to be non-uniformly distributed along the length of all pseudo-chromosomes. Indeed, strong purifying selection against deleterious insertions of TE in gene-rich regions of high recombination has been one of the proposed models to explain heterogeneity in TE distribution along chromosomes (Bartolomé et al. 2002). Consistent with this expectation, heterochromatic pericentromeric regions, marked by suppression of recombination, had consistently low GC content, were gene-poor and enriched with high copy numbers of TEs as compared to the GC-rich euchromatic region. Furthermore, the relationship between STRs and transposable elements has been subject to debate. Preferential accumulation of STRs outside TE-rich regions has been documented in several species (Morgante et al. 2002; Guo et al. 2009). However, in some species this negative correlation has been shown to be specific to certain chromosomes (Li et al. 2017). In our data, we consistently found a significantly lower proportion of STRs in the pericentromeric regions, and the negative correlation of the STRs with TEs held true for all individual pseudo-chromosomes. Further, the locations of the identified centromere-associated repeat were closely associated with the regions of reduced recombination rates. We also found a consistent relative enrichment of LTR elements (as compared to DNA elements) in the pericentromeric regions in both the nine- and three-spined stickleback genomes. The tendency of retrotransposon accumulation in centromeric regions and their potential contribution to the evolution of centromeres has been described earlier in many species (Kent et al. 2017). Further, chromatin immunoprecipitation sequencing along with fluorescence in situ hybridization could be useful in confirming the sequence composition and abundance of these centromere-specific repeats, as well as to gain insights into the repeat organization in functional centromeres of the nine-spined stickleback.

We estimated the age of divergence between the three- and nine-spined sticklebacks to be 26.6 MYA. The estimate is consistent with fossil evidence showing that the three-spined stickleback had diverged from nine-spined stickleback, and also from its congener, the blackspotted stickleback (*G*. *wheatlandi*), at least ~13 MYA (Bell et al. 2009). Our age estimate is considerably larger than a previous estimate that placed the same divergence at 15.86 MYA based on inferred substitution rates for mitochondrial DNA (Aldenhoven et al. 2010) and has been commonly used to as a calibration for phylogenetic analyses in sticklebacks (e.g.(Teacher et al. 2011; Nelson and Cresko 2018)). However, the accuracy of divergence time estimates from mitochondrial DNA has generally been questioned (Brandley et al. 2011; Zheng et al. 2011; Mulcahy et al. 2012). Further, supporting our estimates is a recent, likelihood-based phylogeny, including representatives of Gasterosteiform species (Rabosky et al. 2018). In this study, the divergence of nine-spined and three-spined sticklebacks was estimated to be 29.6 MYA, using nuclear markers. Further, the phylogeny also dates the divergence of *G. wheatlandi*, closer to three-spined stickleback, at 14.3 MYA, consistent with the 13 MYA oldest three-spine stickleback fossil. Considering that the relative age of nine-spined stickleback to that of the *G. wheatlandi* is about twice the age from the phylogeny inferred in Rabosky et al. (Rabosky et al. 2018)), the age of the nine-spined stickleback divergence is expected to be at least on the order of 25-30 MYA. Thus, our estimate, being fairly robust to different filtering methods applied, supports that the divergence time between the nine-spined and three-spined sticklebacks occurred earlier than previously believed.

Rearrangements involving chromosomal inversions and translocation are believed to drive speciation by creating reproductive isolation (Faria and Navarro 2010). The rapid evolution of stickleback karyotypes, as apparent from the diversity in the number of chromosomes and chromosome arms, as well as the sex chromosome systems, has attracted considerable interest (- R. Chen and Reisman 1970; Ross et al. 2009; Ocalewicz et al. 2011; Urton et al. 2011). Based on the pericentromeric regions defined using the centromere-associated repeats and reduced recombination rates, we inspected the pericentromeric rearrangements reported between three-and four-spined sticklebacks, and investigated rearrangements potentially implicated in the difference in chromosomal morphologies between nine-spined and three-spined sticklebacks. Nine-spined stickleback has the highest number of chromosome arms among the sticklebacks (Ocalewicz et al. 2011). Pericentric inversions around the centromere often drive such increases in NF. Using the karyotype derived from the assembled pseudo-chromosomes, we could highlight pericentric inversion events that potentially are responsible for some of the differences in chromosomal morphologies between three-spined and nine-spined sticklebacks.

The diversity in the hemoglobin gene family has been well studied in many vertebrate species. In teleost fish, hemoglobin genes often occur in two distinct unlinked clusters, called MN and LA clusters, located on different chromosomes (Opazo et al. 2013). In three-spined stickleback, a high internal similarity between genes in the MN cluster, potentially harboring a recent *en bloc* duplication comprising of Hba-Hbb or Hba-Hbb-Hba-Hbb has been suggested earlier (Opazo et al. 2013). Interestingly, we identified a recent duplication in the MN hemoglobin cluster, leading to a total of 15 hemoglobin alpha and beta genes in the nine-spined stickleback assembly, in contrast to 11 annotated hemoglobin genes in the MN cluster of the three-spined stickleback assembly. Multiplicity of hemoglobin genes has often been implicated in the ability of tolerating a wide range of environmental stressors (Borza et al. 2009; Opazo et al. 2013). The observed difference in the hemoglobin cluster between three- and nine-spined sticklebacks could thus be of potential biological interest if it is associated with the differing ability to tolerate lower oxygen levels in the two species (Lewis et al. 1972). However, the possibility that this difference in gene copy numbers in the two species could partly stem from the differences in the quality of assemblies cannot be excluded. The long-read data in our assembly potentially resolved some of the highly identical regions arising from very recent duplications, which might not be represented in three-spined stickleback genome assembly. Our analysis of the MN cluster in nine-spined stickleback showed a larger region of high internal similarity and populated by repetitive elements. The duplication could thus be a result of non-allelic homologous recombination owing to the presence of repetitive elements. Highly similar regions of segmental duplications are also often a source of genomic rearrangements due to high frequency of possible misalignments (Stankiewicz and Lupski 2002). The higher tendency of rearrangements in such regions is linked with high occurrences of copy number variations (CNVs) (Perry et al. 2008). In line with this, our analyses of population samples revealed extensive copy number variation in hemoglobin genes even between closely-related populations, suggesting that duplications of individual hemoglobin genes may be of frequent occurrence in the nine-spined sticklebacks inhabiting small water bodies with varying oxygen levels.

## Conclusions

Chromosome-scale whole genome assemblies are a critical resource for elucidating genomic underpinnings and evolutionary forces driving variation in genome structure and organization among different species. With the advent of long-read sequencing technologies, complex genomes have been assembled to a fairly high-quality owing to improved resolution of complex and repetitive regions. The new chromosome-scale assembly of the nine-spined stickleback genome, including detailed analyses of repetitive regions, provides a valuable resource to comparative genomic studies, as well as a solid template for population genomic studies of stickleback fishes. Furthermore, new phylogenetic analyses based on large number of protein coding genes support the notion that the divergence of three- and nine-spined sticklebacks took place about 26 MYA, that is, much earlier than the minimum estimate suggested by the fossil record. Finally, the results regarding extensive copy-number variation in haemoglobin genes, even among populations diverged less than 8000 years ago, suggesting that these genes will deserve further attention in studies seeking to understand the genetic underpinnings of local adaptation in stickleback fishes.

## Methods

### Sampling, DNA extraction and Sequencing

The sequenced male individual was caught April 28 2014 from Pyöreälampi pond from northwestern Finland (66°15’’N; 29°26’’E). This small (< 5 ha surface area) isolated pond was selected as the source because the level of genetic variability in this pond is very low, as revealed by earlier population genetic studies (Shikano et al. 2010). Genomic DNA was extracted from muscle tissue using phenol-chloroform method and fragmented to 20 kb size. All libraries were size selected using BluePippin (4-7 kb) and sequenced on PacBio RSII in a total of 86 SMRT cells (63 SMRT cells with P6v1/C4 chemistry, 23 SMRT cells with P6v2/C4 chemistry). For short-read sequencing, paired-end sequencing Illumina HiSeq 2000 (rapid run 2×250 nt) was performed for the same individual.

### Linkage map construction

Three F_2_-generation interpopulation crosses between pond and marine nine-spined sticklebacks were used as linkage mapping populations. Each of them consisted of first crossing an adult marine nine-spined stickleback female from Southern Finland (Helsinki, 60°13′N, 25°11′E) to a pond male from three different populations (*viz.* Rytilampi, Finland, 66°23′N, 29°19′E; Pyöreälampi (Finland) 66°15′N, 29°26′E and Bynästjärnen (Sweden) 64°27’N;19°26’E in 2006, 2011 and 2012, respectively. After the F_1_ generations fish had matured, F_2_ generations were created by single full-sib mating within each hybrid cross. Fish in both parental generations and the resulting F_2_ offspring in the three crosses were RAD sequenced as described earlier for two of the crosses in (Rastas et al. 2015).

The linkage mapping followed the Lep-MAP3 (LM3) pipeline (Rastas 2017). First, the individual fastq files were mapped to the contig assembly using bwa mem (v 0.7.10) (Li 2013) and sorted bam files were created using samtools (v 1.3.1) (Li et al. 2009). Second, the LM3 pipeline (samtools mpileup and custom scripts) was used to produce proper data for mapping, following with ParentCall2 module with XLimit=2 parameter to call markers from autosomes and the sex chromosome. Third, Filtering2 module was run on the data with dataTolerance=0.001 parameter to remove markers segregating in non-Mendelian fashion (1:1000 by chance). SeparateChromosomes2 was then run with lodLimit=75 finding 21 (major) linkage groups. These group names were mapped to chromosome names used in (Rastas et al., GBE 2015). After this, JoinSingles2All was run with lodLimit=60 and lodDifference=10 to add more markers into linkage groups. After these steps the maps had over 89,000 markers assigned to 21 chromosomes.

In the next step, the OrderMarker2 module was run on each chromosome twice with parameters informativeMask=13 and 23, removing markers informative only for the female or male parent, respectively. The reason for constructing two maps was to reduce the uncertainty in map position caused by markers informative only for one but different parent (having no direct information between each other). The orders were inspected with the LMPlot module and Marey maps were made using custom R scripts.

### Genome Assembly

The raw reads were error-corrected using the hierarchical genome assembly process (HGAP) and assembled using Celera assembler. Genome assembly was performed using Celera assembler 8.2, followed by polishing using Quiver (Chin et al. 2013), yielding 5,303 scaffolds with a total size of 522 Mbp. Quality checked Illumina HiSeq2500 reads were then mapped to the contigs using BWA-MEM (v0.7.10) (Li 2013) and the alignment was used to polish the PacBio assembly using Pilon (v 1.9) (Walker et al. 2014). Validation of the assembly in terms of its completeness was performed by searching for core eukaryotic orthologous genes using BUSCO (v3.0.1) (Simão et al. 2015). The contigs consisting of mitochondrial genome sequences were discarded owing to misassembly, and we then used the mitochondrial sequence described in Guo et al. (Guo et al. 2016) as the reference sequence. The Illumina reads mapping to the mitochondrial genome were then extracted and variants were inferred using samtools (Li et al. 2009) mpileup. A consensus mitochondrial genome sequence was generated using GATK (DePristo et al. 2011) FastaAlternateReferenceMaker and added to the assembly.

### Anchoring contigs to the linkage map

Each marker in the linkage map had a coordinate in the contig assembly, and this information could be used directly to anchor contigs into pseudo-chromosomes. Each contig was placed on most abundant pseudo-chromosome among linkage map markers in it. For the contigs where multiple SNPs supported different linkage groups, based on the number of such matches, the contig was either broken or assigned to the linkage group with largest number of hits. The median map position for each contig was computed and it was used to approximate place contigs within pseudo-chromosomes. A gap of 200 bases was inserted in between the anchored contigs. Contigs with only one marker were not anchored, except for the X chromosome region where typical contig lengths were shorter. The exact location and orientation of contigs and chimeric contigs were further fixed by manually inspecting the Marey maps. The recombination rate was estimated as the derivative of a non-decreasing spline function fitted to the Marey map using module cobs (He and Ng 1999) in R. The R code to estimate recombination rate is included in the supplement.

### Transcriptome assembly

The RNA-seq data from brain and liver for four individuals were generated using Illumina paired-end sequencing. The read data were evaluated for quality using FastQC and pooled to assemble using Trinity package (v2.0.6) (Grabherr et al. 2011; Emms and Kelly 2015). *In silico* normalization was performed on the reads prior to the assembly (Haas et al. 2013). Using default parameters, the Trinity *de novo* pipeline resulted in 255,469 transcripts with a CEGMA completeness of around 89% (complete match) to 100% (partial match). Transcript abundance estimates were obtained using the RSEM method implemented within the Trinity package. The assembled transcripts were then filtered based on a FPKM (fragments per kilobase of transcript per million fragments sequenced) threshold of 0.05, resulting in 123,174 transcripts.

### Repeat annotation

Repetitive sequences in the *P. pungitius* genome assembly were identified using both *de novo* and homology methods. The de novo repeat identification was performed using Repeat Modeler (v1.0.8, http://www.repeatmasker.org/RepeatModeler) and Transposon PSI (http://transposonpsi.sourceforge.net/). The sequences were combined and clustered using USEARCH (v9.2.64) (Edgar 2010) at a threshold of 80%. Additionally, full length LTR sequences were identified using LTR_finder (Xu and Wang 2007) and LTRharvest (Ellinghaus et al. 2008), and were combined using LTR_retriver (Ou and Jiang 2018). The sequences were classified using RepeatClassifier (a part of Repeatmodeler package), TEclass, Censor, and Dfam database (Wheeler et al. 2013). RepeatMasker (v 4.0.7) was used to annotate the identified repeat elements on the assembly.

### Genome Annotation

The annotation was done on the repeat-masked genome following a two-pass approach using MAKER2 (v 2.31.9) pipeline (Holt and Yandell 2011). The first round used Genemark-ES (v 2.3e) (Lomsadze et al. 2005) for *ab initio* prediction of genes and SNAP model trained on CEGMA genes. The *de novo* transcriptome assembly, UniProt/SwissProt database (UniProt Consortium 2015) and *G. aculeatus* CDS sequences were used as evidence sets for the prediction of gene models. For the second round, SNAP (v 20131129) (Korf 2004) and AUGUSTUS (v 3.2.2) (Stanke et al. 2008) were trained on the gene model predicted from the first pass. Functional annotation was performed using BLASTP against UniProt proteins with an E-value threshold of 1e-5, and InterProScan (v 5.4-47) (Jones et al. 2014) was used for domain annotation. The resulting gene models were filtered to retain those with ‘annotation edit distance’ (AED) value of 0.5 or less, having PFAM annotations and significant hits to known proteins against Uniprot DB (e-value 1e-5).

### Tandem repeat analysis

To determine the sequence and structure centromeric repeat sequence, a random sample of 500,000 PacBio subreads were extracted. These were processed to retained only sequences with length greater than 1,000 bp and less than 5% Ns, and low complexity regions were masked using DUST filter as done in (Melters et al. 2013). Tandem repeat finder (v4.0.7) (Benson 1999) was run on the resulting sequences. Sequences greater than 100 bp and occupying more than 80% of the read length were retained as putative centromeric repeats. The most abundant representative repeat sequence of 178 bp length was then aligned to the centromeric repeat sequence in the three-spined stickleback (GacCEN, accession KT321856 (Cech and Peichel 2015) using PRANK (Löytynoja and Goldman 2005). To survey STRs in the genome, Phobos (verison 3.3.12) was run to determine repeats upto unit size of 6, with otherwise default settings. The telomeric repeat arrays were identified by filtering Phobos tandem repeat output to include greater than 10 copies of ‘AACCCT’ repeat.

### Identification of orthogroups and phylogenetic analysis

The protein sequences from zebrafish (GRCz10, Ensembl release 8), Atlantic cod (gadMor2, (Tørresen et al. 2017)), platyfish (Xipmac4.4.2, Ensembl release 89), Nile tilapia (GCF_001858045.1_ASM185804v2), medaka (MEDAKA1, Ensembl release 89), tetraodon (TETRAODON 8.0, Ensembl release 89), fugu (FUGU5), three-spined stickleback (Glazer et al. 2015) and nine-spined stickleback were used for orthologous group analysis using OrthoFinder (v1.0.6) (Emms and Kelly 2015). Clustering into orthologous gene families was done based on best reciprocal blast hits resulting from an all-vs-all blast with E-value threshold of 1e-5.

To perform phylogenetic analysis, single copy orthologs were extracted and further filtered to only retain complete BUSCO proteins (based on BUSCO Actinopterygii odb9). The protein alignments were then converted to codon alignment using pal2nal (Suyama et al. 2006), trimmed using gblocks and further filtered to remove the third codon position. The output of pal2nal for these genes was then concatenated and trimmed using gblocks and filtered to retain only the 1st and 2nd codon positions. Firstly, BEAST analysis (version 2.5) was done (Bouckaert et al. 2018) using bmodelTest (Bouckaert and Drummond 2017) and relaxed clock model. The Yule tree model was used with MRCA age calibrations added according to the divergence estimates in Betancur-R et al. (2013) for all the nodes except the sticklebacks and divergence time was estimated for the sticklebacks. The Markov-chain Monte Carlo (MCMC) analysis was run using a chain length of 10 million logging every 10,000th step. Tracer (v1.6.0) (Rambaut et al. 2018) was used to verify that the effective sample size is above 200 for all model parameters, indicating convergence of the MCMC analysis. TreeAnnotator (v2.4.8) was then used to generate a maximum clade credibility tree with median node heights.

To further check the robustness of the obtained estimates, we applied rigourous filters on the orthogroups. For this, BEAST analysis with the same parameters as mentioned above was performed on all the individual 2,691 gene sets. From the resulting trees, all those that did not support monophyly of the two stickleback species were eliminated. Secondly, to eliminate potential misalignments and paralogy, the protein alignments were inspected with a set of stringent thresholds, to remove alignments with outlier-like values of bit score, alignment length and gap open. This set of orthogroups were further filtered based on clock-likeness by excluding alignments for which a high standard deviation of the uncorrelated lognormal (UCLN) (Drummond et al. 2006) molecular clock was inferred in the single-gene BEAST analyses (a threshold of 0.1 for UCLDstddev was chosen). The remaining 778 gene alignments were concatenated to form a supermatrix and BEAST analysis was then rerun with this supermatrix using similar settings as above. MCMC analysis was performed using chain length of 1 billion, logging every 1000 steps with a burn-in of ~25%. The run was monitored using Tracer (v1.6.0) (Rambaut et al. 2018) and TreeAnnotator (v2.4.8) was then used to generate maximum clade credibility tree. The tree resulting from the concatenated alignments of 778 genes was used as an input to gene family analysis using CAFE (v 3) (Han et al. 2013). Cafeerror was run for error model estimation and then CAFE was run with a global lambda estimation using the error model.

### Hemoglobin gene cluster analysis and population analysis

Hemoglobin genes from three-spined stickleback and zebrafish were used to query nine-spined stickleback genome assembly using TBLASTN. The coordinates of the obtained hits were used to extract sequences corresponding to the MN and LA cluster in the assembly. To predict genes in this region, GENSCAN was used with the human model. The predicted protein sequences were aligned using MUSCLE and maximum likelihood trees were generated in MEGA. To calculate read depth across the region, the Illumina reads were first mapped using bwa-mem, bedtools was then used to calculate coverage per base.

Five samples from Pyöreälampi, Finland, and Levin Navolok Bay, Russia, were sequenced to 10X coverage (BGI Genomics, People’s Republic of China). Reads were mapped to the reference genome with bwa mem (v. 0.7.15) and realigned around gaps with GATK IndelRealigner (v. 3.7). Duplicate reads were marked and site-wise sequencing coverage computed with samtools (v. 1.9) markdup and depth, respectively. From these coverage counts, the mean coverage was computed for different repeat element classes, for all coding exons (totalling 205,048 after excluding LG12 which is the sex chromosome) and for the individual genes (coding exons only) within the MN hemoglobin cluster. The coverage for the repeat element classes and the hemoglobin genes were normalised by the mean sequencing coverage over all coding exons.

### Synteny analysis

Large scale gene order synteny between *G. aculeatus* and *P. pungitius* was identified using MCScanX (v1). The collinear blocks with conserved gene order were identified using a BLASTP with e-value 1e-5 and match size of 10. The same was repeated using the Medaka (MEDAKA1, Ensembl release 89) assembly.

## Acknowledgements

We thank the Oulanka Biological Station, and Pia Saarinen in particular, for help in obtaining the individual used to generate the genome assembly. Kirsi Kähkönen and Ave Tooming-Klunderud are thanked for their help with DNA-extractions and sequencing. PacBio sequencing was performed at the Norwegian Sequencing Centre (NSC) and sequencing (Illumina) of population samples at BGI Hong Kong and Institute of Biotechnology, University of Helsinki. Our research was supported by grants from the Academy of Finland (250435, 263722, 265211 and 1307943 to JM), Helsinki Institute for Life Sciences (HiLIFE; H9701-11-109105 to JM), and Marie Curie Intra-European Fellowship within the 7th European Community Framework Programme under REA grant agreement PIEF-GA-2013-624073 (FC & JM). MM was supported by the Norwegian Research Council (FRIPRO). SV and KSJ were supported by UiO through a strategic research initiative.

