## Supplementary File for "Genome sequencing of the nine-spined stickleback (*Pungitius pungitius*) provides insights into chromosome evolution"

### Supplementary Figures

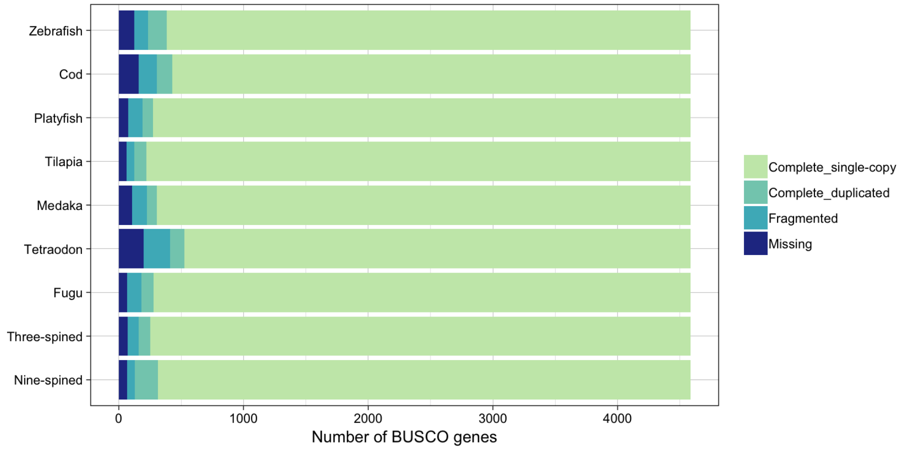


**Supplementary Figure S1:** The BUSCO scores of the nine-spined stickleback genome assembly compared to that of zebrafish (GRCz10, Ensembl release 8), Atlantic cod (gadMor2 (Tørresen et al. 2017)), platyfish (Xipmac4.4.2, Ensembl release 89 ), Nile tilapia (GCF_001858045.1_ASM185804v2), medaka (MEDAKA1, Ensembl release 89), tetraodon (TETRAODON 8.0, Ensembl release 89), fugu (FUGU5) and three-spined stickleback ((Glazer et al. 2015)).

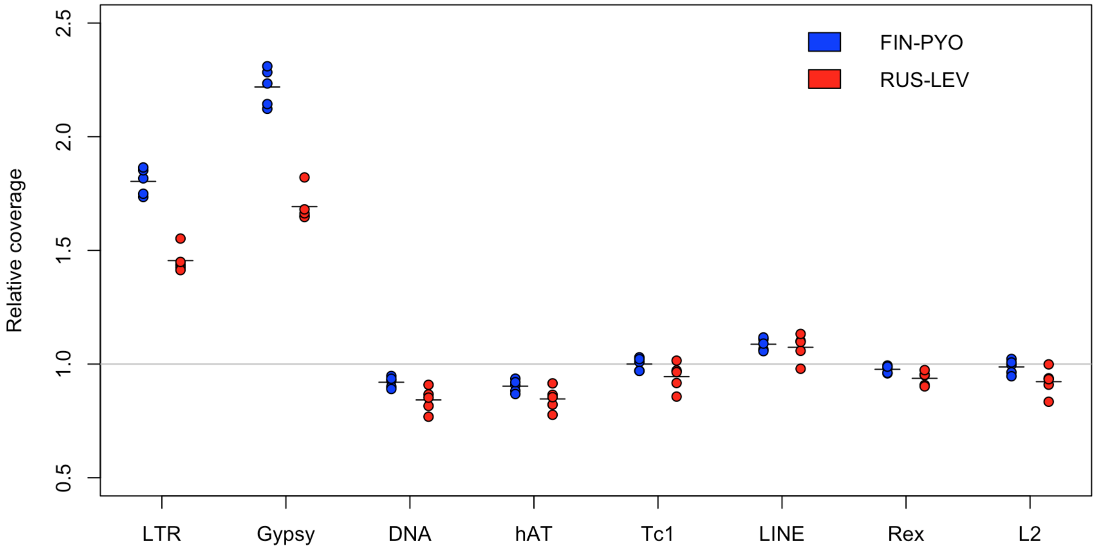


**Supplementary Figure S2:** Each dot represents the mean relative coverage for each of the repeat families and the most abundant sub-families for each of the five individuals sequenced from the two populations, namely, pond population from Pyöreälampi, Finland (FIN-PYO), and from the ancestral marine population in Levin Navolok Bay, Russia (RUS-LEV).

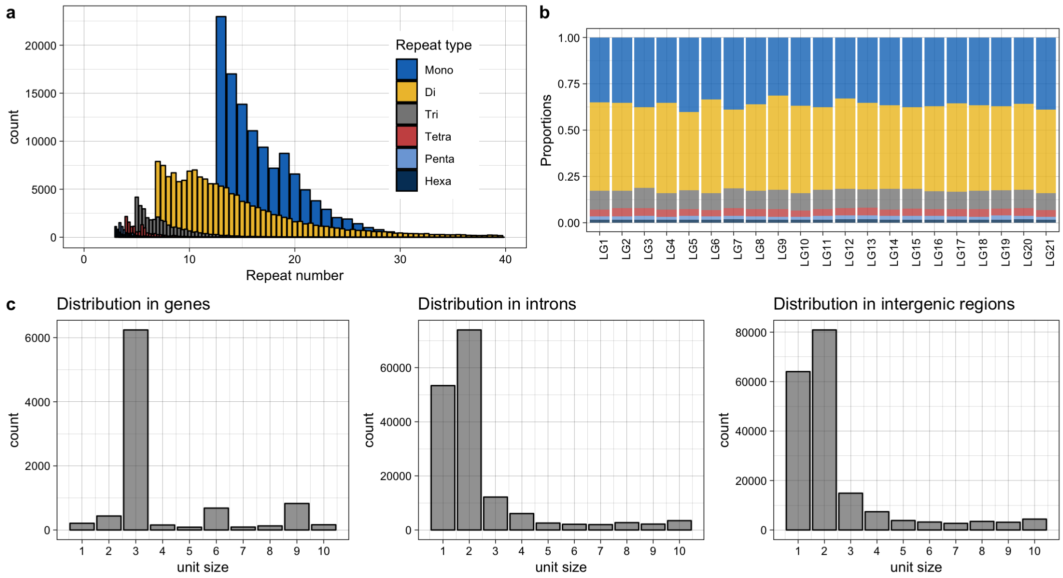


**Supplementary Figure S3**: **a.** Overall distribution of short tandem repeats (STRs) of various unit sizes and number of bases in repeat numbers. **b.** Proportion of each type of STRs across the nine-spined stickleback pseudo-chromosomes. **c-e.** Distribution of STR units in genes, introns and intergenic regions.

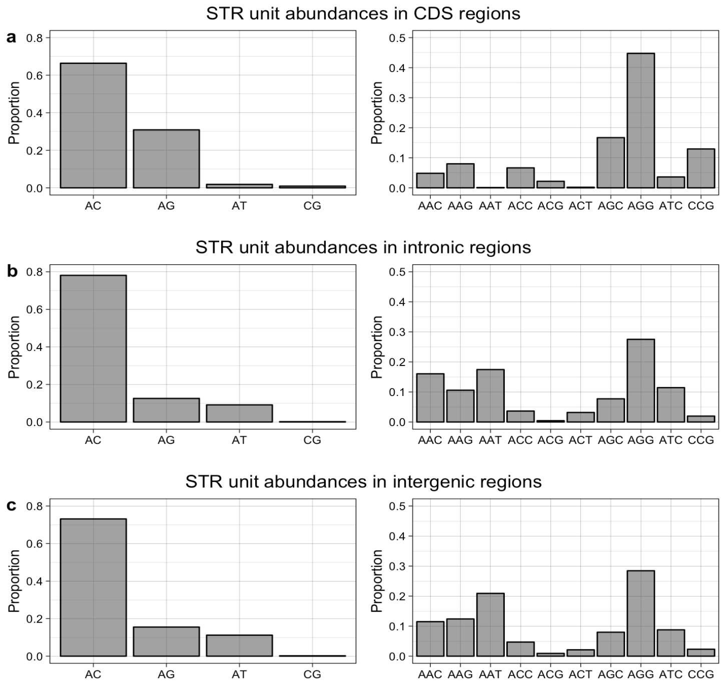


**Supplementary Figure S4**: **a-c** Relative proportions of various di- (left panel) and tri-nucleotide (right panel) repeat motifs in coding, intronic and intergenic regions of the nine-spined stickleback genome respectively.

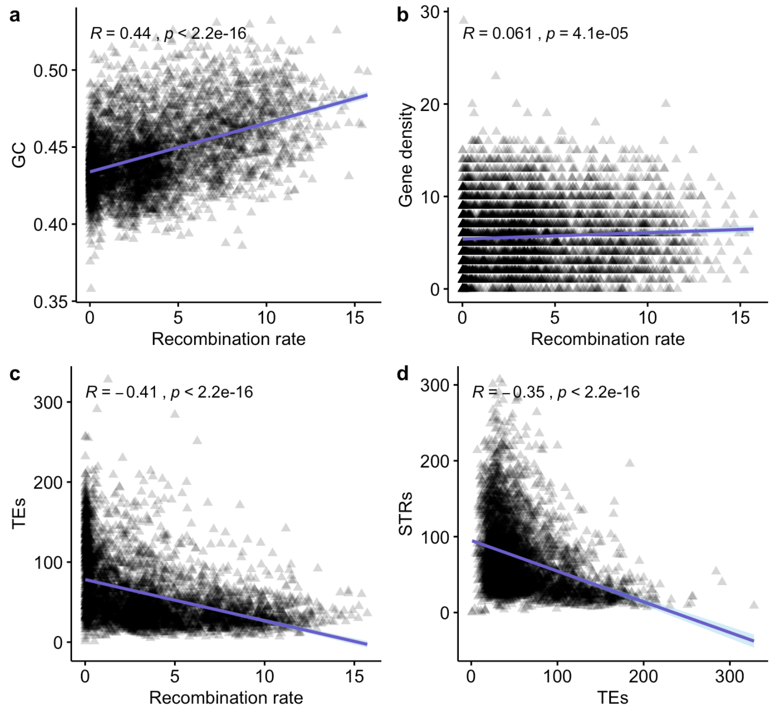


**Supplementary Figure S5:** Global correlations between recombination rate (x-axis in a-c) and (a) GC content, (b) Gene density and (c) Transposable element (TE) density, in 100Mb bins, across all the chromosomes. (d) Global correlations between the counts of TEs and STRs, in 100 Mb bins, across all chromosomes.

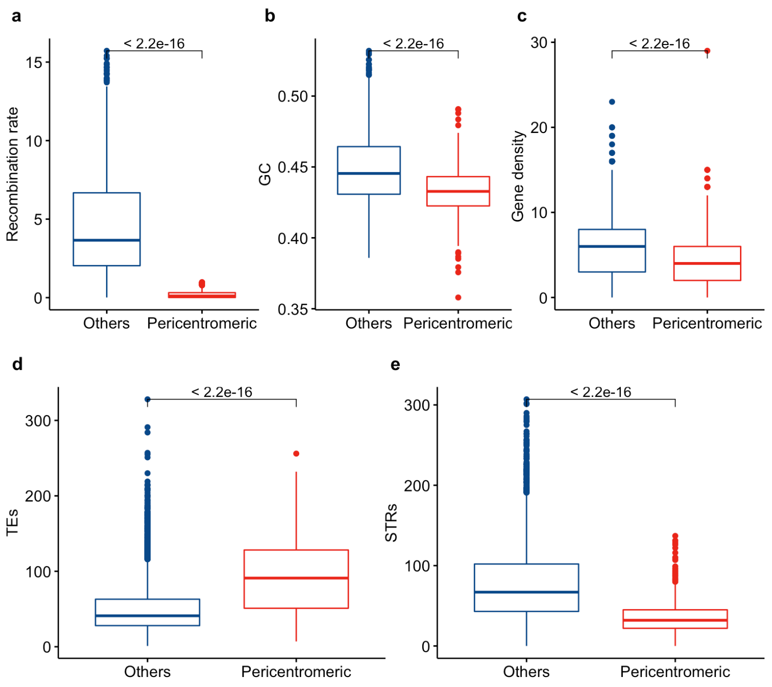


**Supplementary Figure S6:** Distribution of (a) recombination rate, (b) GC content, (c) Gene density, (d) Transposable elements (TEs) and (e) short tandem repeats (STR) densities within and outside the inferred pericentromeric regions.

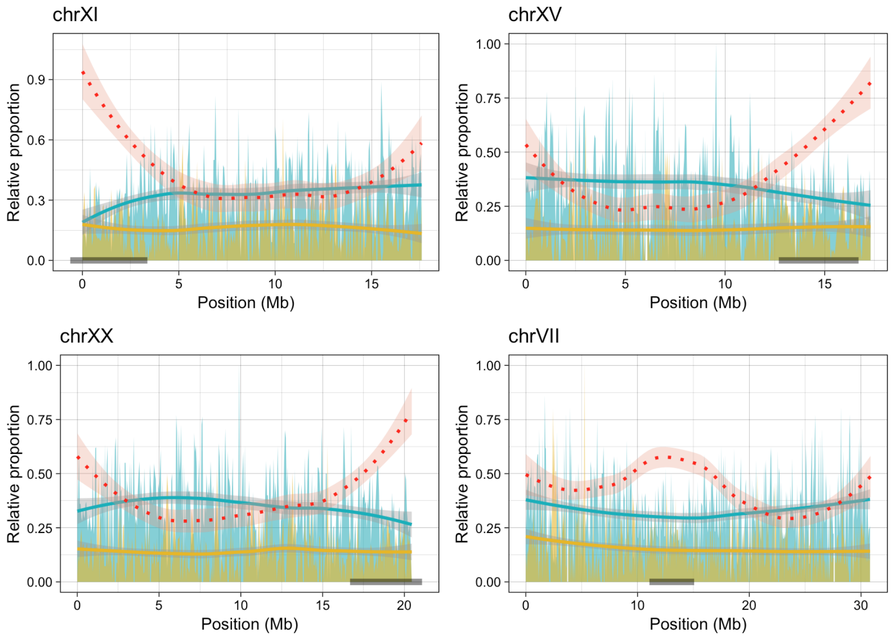


**Supplementary Figure S7:** Distribution of relative proportions of LTR (yellow) and DNA (blue) transposable elements to total repeat content per 50Kb bin along the chromosomes in the three-spined stickleback**.** The red dotted line represents log10 of absolute abundance of LTR-gypsy elements across the chromosomes. The grey rectangle represents the probable pericentromeric region

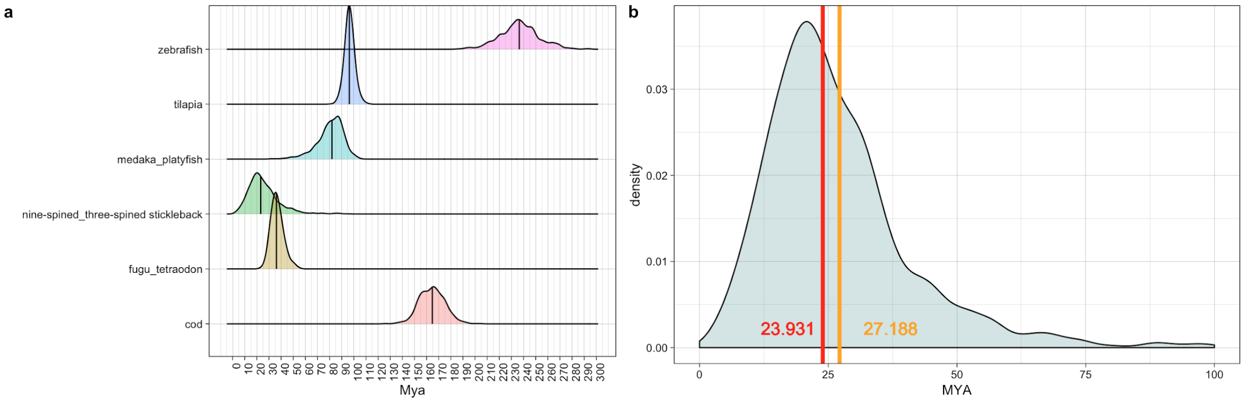


**Supplementary Figure S8:** (a) Divergence time estimates from BEAST runs of 778 individual gene trees and b) distribution of divergences times for the nine-spined stickleback when constraints are placed only on two nodes (zebrafish and Atlantic cod). The red and yellow lines represent the median and mean divergence times respectively.

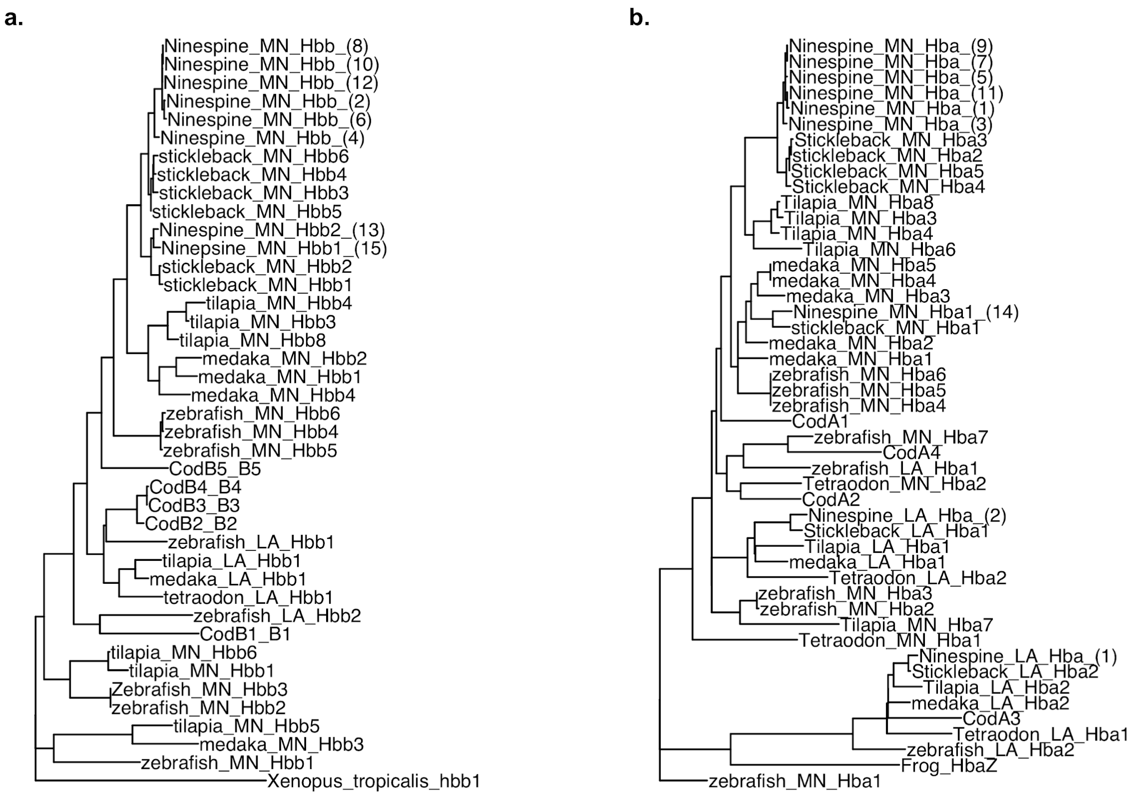


**Supplementary Figure S9:** Phylogenetic tree of protein sequences of Hemoglobin beta (a) and alpha (b).

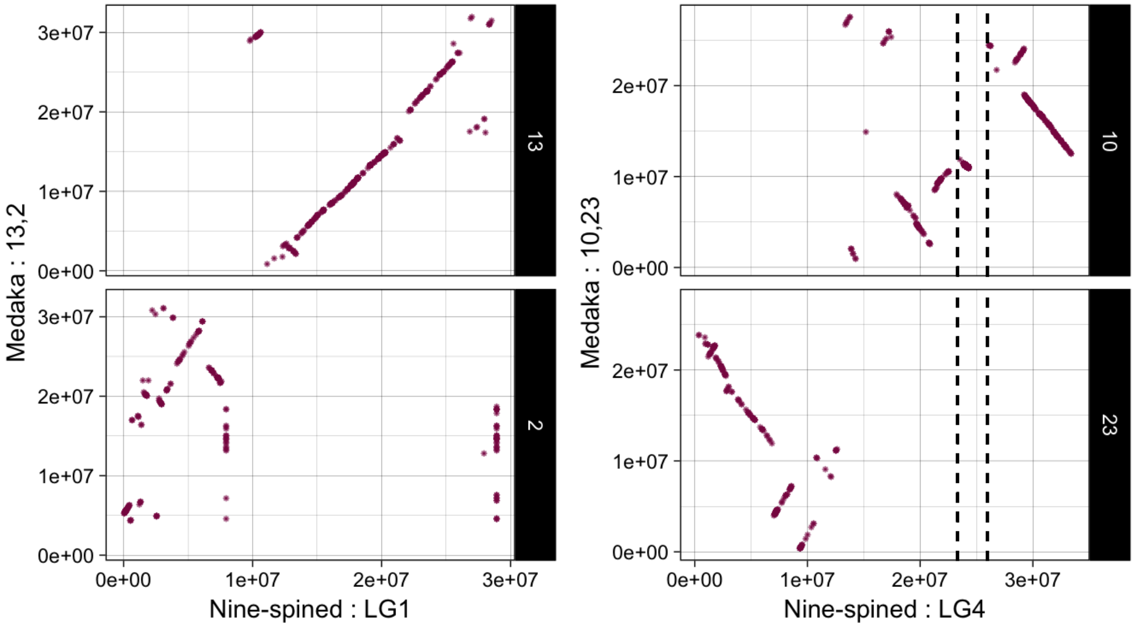


**Supplementary Figure S10: Synteny of nine-spined stickleback LG1 and LG4 with corresponding orthologs in medaka.** The nine-spined stickleback chromosome positions are shown on the x-axis while the y-axis axis represents the corresponding orthologous chromosomes in medaka.

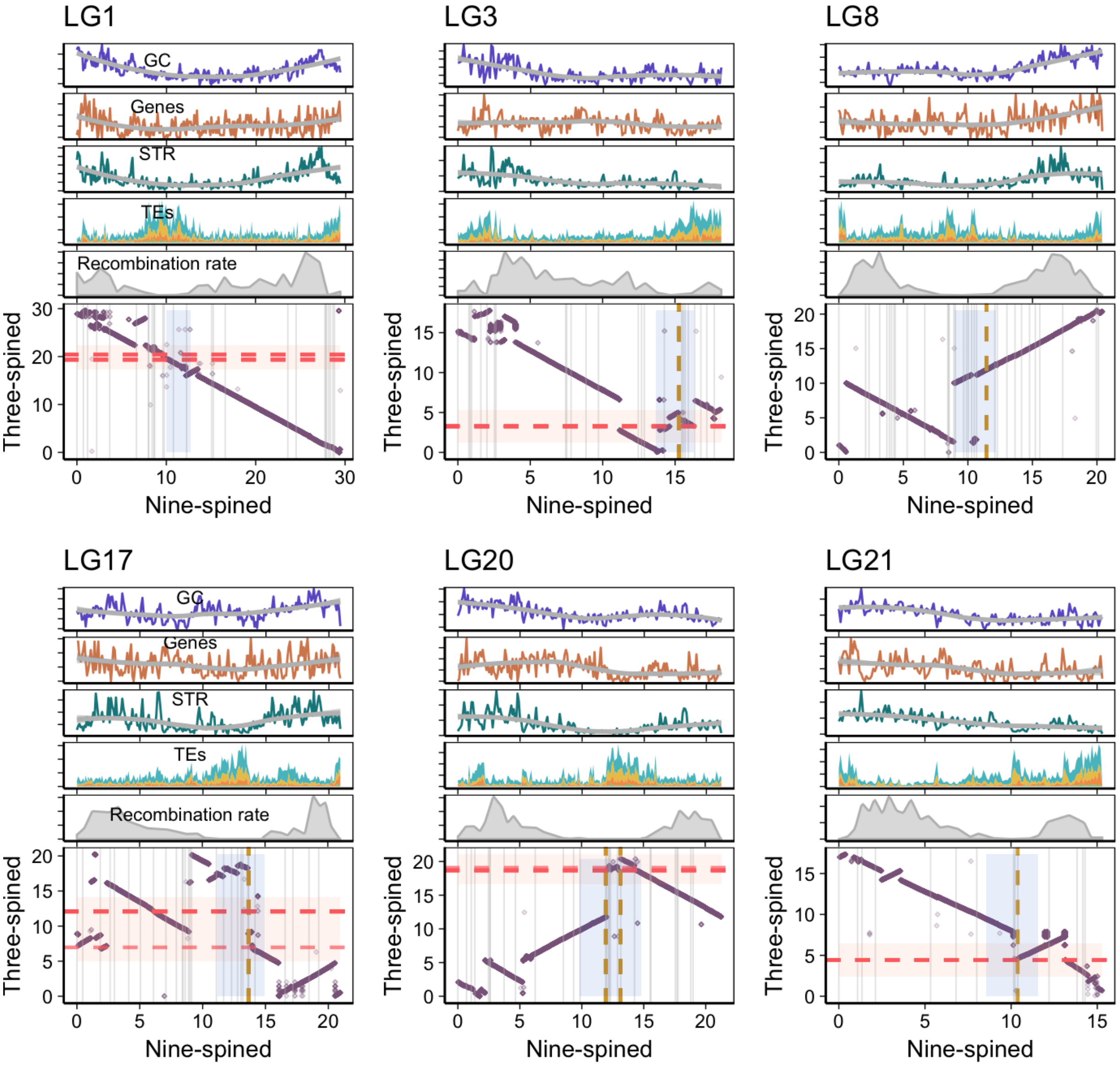


**Supplementary Figure S11**: **Conserved synteny between three-spined and nine-spined stickleback for LG 1, 3, 8, 17, 20 and 21**. Top: The distribution (per 100kb bins) of GC content, gene density, TE density (yellow: LTR, blue: DNA, orange: LINE, green: SINE) and recombination rate along each chromosome. Bottom: Alignment of nine-spined stickleback with the corresponding orthologous three-spined stickleback chromosome. The dotted lines represent the location of the centromeric repeats and the shaded areas represent putative pericentromeric region in the two genomes.

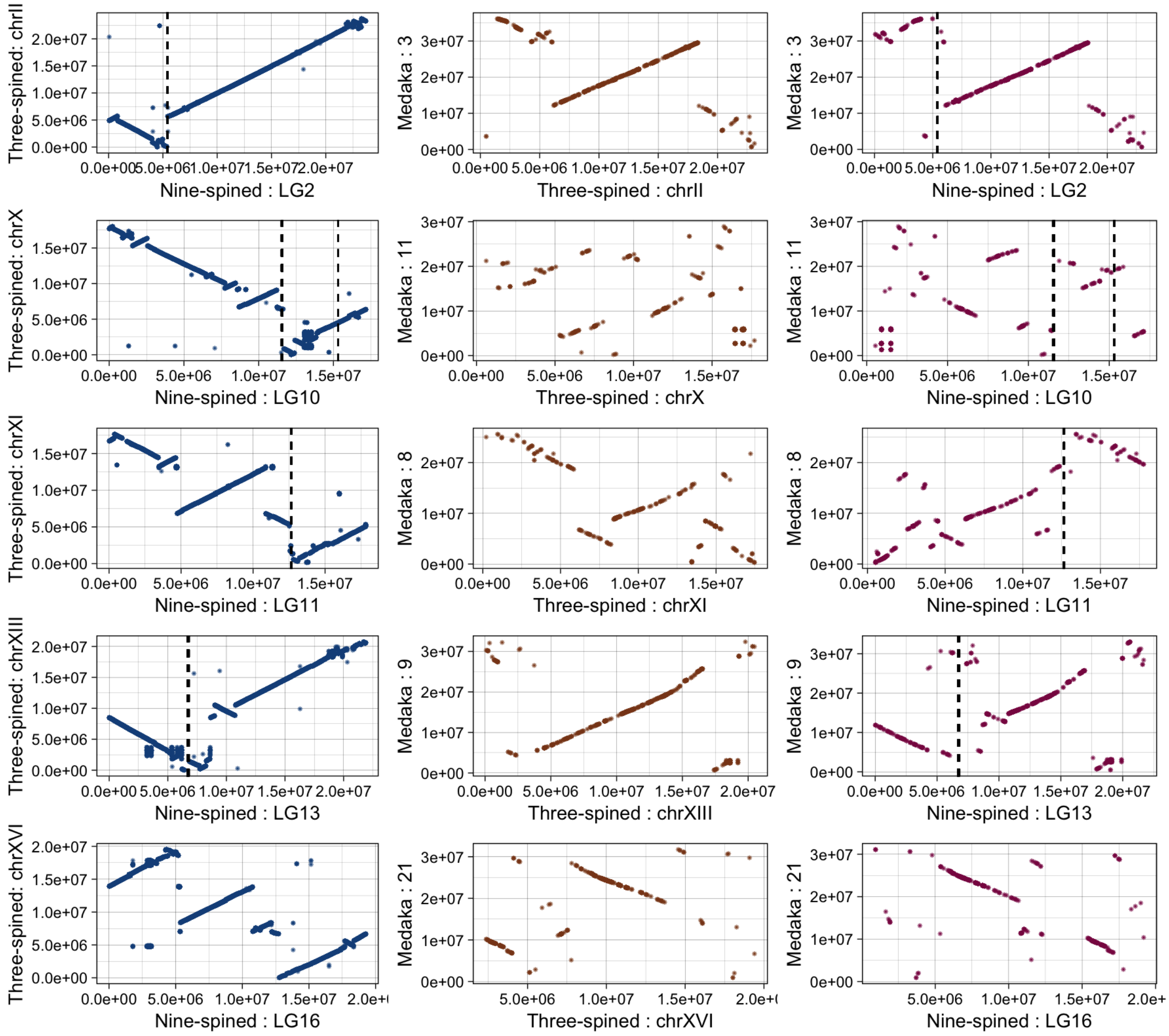


**Supplementary Figure S12**: **Distribution of conserved synteny among the genomes of medaka, nine-spined and three-spined sticklebacks (for nine spined stickleback LG 2, 10, 11, 13 and 16).** First panel in each row represents alignment of a nine-spined stickleback chromosome with three-spined stickleback chromosome, second panel shows three-spined stickleback aligned to medaka chromosome, and the third panel represents nine-spined stickleback chromosome aligned to the corresponding medaka ortholog. Dashed lines represent the location of centromeric tandem repeat in nine-spined stickleback.
